# Accelerated phenology fails to buffer fitness loss from delayed rain onset in a clade of wildflowers

**DOI:** 10.1101/2024.06.12.598693

**Authors:** Samantha J. Worthy, Sarah R. Ashlock, Arquel Miller, Julin N. Maloof, Sharon Y. Strauss, Jennifer R. Gremer, Johanna Schmitt

## Abstract

The timing of early life cycle events has cascading effects on phenology and fitness. These effects may be critical for climate resilience of plant populations, especially in Mediterranean environments, where delayed rainfall onset causes delayed germination. To examine impacts of germination timing on ten species of the *Streptanthus*/*Caulanthus* clade, we induced germination across a range of dates in ambient seasonal conditions and recorded phenological and fitness traits. Later germination cohorts accelerated flowering, partially stabilizing flowering date, but the degree of this compensatory plasticity differed across species. Fitness declined with later germination; the magnitude of this decline depended on the balance between direct negative effects of later germination and compensatory positive effects of accelerated flowering. The resulting species’ differences in fitness responses suggest differential vulnerability to climate change. Species from wetter, cooler, less variable habitats accelerated flowering more and declined less in seed set with later germination, suggesting climate adaptation. However, other fitness responses to germination timing, like first year fitness, were evolutionarily labile across the clade and unrelated to climate. Although compensatory phenological plasticity may buffer the impacts of delayed germination, it cannot prevent long term declines in population fitness as fall rains come later with climate change.

## Introduction

The seasonal timing of early life cycle events such as birth, hatching, or seed germination has critical effects on fitness in many organisms (Kalisz 1986; Donohue 2002, 2005; Donohue et al. 2005; van Asch and Visser 2007). The timing of these events determines the effective length of the growing season and the environmental conditions organisms experience throughout the life cycle, including environmental cues mediating the timing of later life cycle events such as reproduction (Galloway 2001; Donohue 2005; Donohue et al. 2005; Wilczek et al. 2009). However, climate change is altering the timing and length of the growing season, as well as the environmental cues driving seasonal phenology for many organisms (Parmesan and Hanley, 2015; Wadgymar et al. 2018; Bonamour et al. 2019; Bernhardt et al. 2020). An important question is whether phenological plasticity can buffer the population fitness impacts of such environmental change (Ghalambor et al. 2007; Nicotra et al. 2010; Chevin et al. 2010; Duputié et al. 2015; Kingsolver and Buckley 2017; Scheiner et al. 2020; Gauzere et al. 2020; Zettlemoyer et al. 2024). If so, the resilience of species’ fitness to changing climate may depend upon the phenological consequences of early life cycle events (Gremer et al. 2020a; Martínez-Berdeja et al. 2023).

In plants, seed germination timing is a critical early event that determines the seasonal conditions that plants experience throughout the life cycle (Donohue 2005). A species’ germination niche - the range of environmental conditions under which seeds can germinate-determines its germination phenology (Donohue et al. 2010; Martínez-Berdeja et al. 2023; Worthy et al. 2023). In seasonally dry climates, including Mediterranean climates, the timing of precipitation onset relative to seasonal environmental cues regulating germination determines whether and when seedlings will emerge (Kimball et al. 2010; Martínez-Berdeja et al. 2020, 2023; Worthy et al. 2023). Germination timing in turn determines the length of growing season, as well as exposure to environmental cues that mediate the timing of later events such as flowering and seed set. These cascading effects of germination timing may be important to species’ fitness as climate change alters the timing and variability of seasonal precipitation in Mediterranean regions (Swain et al. 2018; Dong et al. 2019; Luković et al. 2021). In California, germination-triggering rains are coming later in fall, with a more compressed rainy season (Luković et al. 2021). This delay in rainfall onset constrains seeds to germinate later in the year, under different seasonal conditions and with a shorter potential growing season (Worthy et al. 2023).

Seasonal delays in germination timing may affect species’ fitness directly, through effects on growing season environment or length, or indirectly through effects on reproductive phenology. Flowering time is often under strong selection, which usually favors earlier flowering (Munguía-Rosas et al. 2011; Anderson et al. 2012; Austen et al. 2017; Ensing et al. 2021; Zettlemoyer et al. 2024). In a shorter potential growing season, such selection may favor accelerated flowering at a smaller size to assure reproduction (Cohen 1976). If so, acceleration of flowering in later germinating cohorts, mediated by seasonal environmental cues such as increasing day length or accumulation of photothermal units (Wilczek et al. 2009), could be a form of compensatory phenotypic plasticity mitigating fitness impacts of shorter growing seasons. On the other hand, if later germination prevents exposure to critical seasonal cues such as accumulation of chilling hours (vernalization), the transition to flowering may be delayed or prevented with critical consequences for life history and fitness (Galloway and Etterson 2007; Gremer et al. 2020a). Species differences in phenological plasticity and fitness resilience to germination timing may contribute to differential vulnerability to climate change (Matesanz et al. 2010; Chevin et al. 2012).

Here, we examine the phenological and fitness consequences of germination timing in the *Streptanthus/Caulanthus* clade (Brassicaceae). This group has diversified from desert origins into a wide range of Mediterranean climates across the California Floristic Province (Axelrod 1958; Cacho et al. 2014, 2021). In a previous experiment, we found evidence that species differ across the clade in responses to delayed rainfall onset at the earliest life history stage (Worthy et al. 2023). Germination declined with later rainfall onset in many species, but the magnitude of response varied significantly across the phylogeny (Worthy et al. 2023). Here, we investigate the potential impacts of germination delays on subsequent flowering phenology and fitness. To this end, we experimentally induced germination at successive intervals through the growing season, thus forcing plants to experience a range of natural seasonal environments both within and beyond their typical germination window. We asked: (1) Does flowering phenology respond to germination timing? Depending upon species’ responses to seasonal cues, flowering in later germination cohorts might be either accelerated (Wilczek et al. 2009; Wu and Owen 2017; Olliff-Yang and Ackerly 2021; Martinez-Berdeja et al. 2023) or delayed (Gremer et al. 2020a). (2) Does germination timing affect fitness, either directly or indirectly through effects on flowering phenology?, (3) How have responses of flowering time and fitness to germination timing diversified across the clade, and do patterns reflect climate of origin?, and (4) Can variation among species in responses to germination timing lead to differential vulnerability to climate change? Understanding variation in responses to germination timing can reveal species’ differences in vulnerability to climate change at later life stages, compounding the potential effects of climate change on germination alone (Walck et al. 2011; Worthy et al. 2023).

## Methods

### Study System

To examine the phenological and fitness consequences of variable germination timing, we used 12 populations from 10 species that span the *Streptanthus* and *Caulanthus* genera of the *Streptanthus* clade of Brassicaceae: *Caulanthus anceps* (CAAN), *C. coulteri* (CACO), *C. inflatus* (CAIN), *Streptanthus breweri* (STBR), *S. diversifolius* (STDI), *S. drepanoides* (STDR), *S. glandulosus* (STGL), *S. insignis* (STIN), *S. polygaloides* (STPO), and *S. tortuosus* (STTO) (Table S1; Cacho et al. 2014). Henceforth, we refer to all species and populations by the abbreviations given above. These species are found across a broad latitudinal range that encompasses the California Floristic Province, spanning a wide range of mean and variability in temperature and precipitation (Cacho et al. 2021; Worthy et al. 2023). They all experience growing seasons dictated by the Mediterranean climate of California where germination-triggering rain events start the growing season in the fall and onset of drought ends the growing season. All species are winter annuals except for STTO, which has variation in life histories within and across populations, with facultative winter annual, biennial, and perennial life histories observed in some populations (Gremer et al. 2020a). The STTO population used in this study (TM2) is largely winter annual but also exhibits biennial and perennial life histories depending on germination timing and exposure to vernalization cues (Gremer et al. 2020b, Gremer unpublished data). Hereafter, we will refer to the STTO life history as facultative biennial.

Eight of the species in this study were represented by one population approximately centrally located within the species’ range-wide climatic space (Figure S1). Two species, CAAN and CAIN, were opportunistically represented by two populations because of seed availability. In all analyses, population was initially included, but was not a significant predictor so results are presented and discussed at the species level.

Seeds for the experiment were collected from the field in 2019 (June – August) or harvested from plants grown out simultaneously in a “screenhouse” common garden (Table S1). The screenhouse has a clear plastic roof and screened walls that allow exposure to ambient temperatures and day length but controlled watering and pollination. Seeds from approximately 20 maternal families were pooled for each species and population. All seeds were stored dry at room temperature in the dark from collection until the start of the experiment.

### Experimental Design

To examine phenological and fitness responses to seasonal germination timing, we induced germination on different dates to create eight staggered seasonal germination timing cohorts, one cohort planted every three to four weeks from September 20, 2021, to March 7, 2022. These cohorts spanned the natural germination timing in the field for these populations, as well as extending timing earlier and later in the season. By forcing seeds to germinate outside their species’ normal seasonal germination niche, we could observe phenology and fitness responses to germination timing across a wider range of seasonal environments, simulating historical extremes of rain onset and future climate change scenarios.

For each germination timing cohort, 75 seeds of each species were sown into individual cells of Landmark 98 germination plug trays with a goal of 24 seedlings for later transplant. Trays were filled with a mix of 2/3 UC Davis potting soil (equal parts sand, compost, and peat moss with dolomite) and ⅓ coarse 16 grit sand, watered until saturation. Seed coats of the *Caulanthus* species were scarified using a scalpel prior to sowing to remove physical dormancy barriers to germination observed in prior unpublished experiments (Worthy et al. unpublished data; LoPresti et al. 2019). After planting, trays were placed into growth chambers (E7/2 Conviron, Winnipeg, Manitoba, Canada) for 17 days with 12-h daylight cycles at varying temperatures to promote germination (Table S2).

For each planting cohort, germination trays were placed in the screenhouse after the 17-day germination period in growth chambers. After a 24-h adjustment period, seedlings were transplanted into individual cone-tainers (164-ml Stuewe and Sons SC10) filled with the same soil mixture as the germination trays. The height of each seedling was recorded to use as a size covariate in analyses, and cones were placed on a mist bench for three weeks to minimize transplant shock. Following this acclimation period, cones were placed randomly into population-specific racks distributed across four screenhouse benches. Sample size varied among populations and among cohorts due to variation in germination ranging from 1 to 24 individuals (Table S1). When plants flowered, we hand-pollinated them every two or three days to ensure pollination. We used cotton swabs to collect pollen from open flowers of at least three individuals per species and then distributed pollen to all flowering conspecifics.

During the study, plants were watered using a drip irrigation system with each species assigned to one of three watering regimes: low (49.50 mL/week), medium (57.75 mL/week), or high (75.25 mL/week). In pilot experiments, we found that desert species got fungal disease with too much water, while species occupying wetter climates suffered under drier desert watering regimes. We therefore grouped species into watering regimes similar to annual rainfall at their field sites, determined by comparing 30-year average annual precipitation (1991-2020) for each populations’ collection site extracted from the PRISM database (PRISM Climate Group 2014; Table S1). These precipitation amounts were converted to mm per week of the study, then to mL (1 mm = 1L per m^2^), and then used to calculate the length of time the irrigation system should run to deliver the desired quantity of water (based on the average flow rate of the drip irrigation system, 16.5 mL per minute). When necessitated by heat waves, additional water was supplied with proportional increases across the watering groups (Table S3). To simulate a natural end to the season, watering was tapered off over a three-week period prior to the end of the study on June 13, 2022.

### Environmental measurements

To quantify seasonal environments experienced by successive germination cohorts, we recorded hourly temperature in the screenhouse with temperature loggers (Thermochron DS1921G iButtons) buried in soil-filled cones from the date of transplant to the end of the study for each germination timing cohort. We used these data to calculate two metrics of cumulative growing season conditions for each germination timing cohort: chill portion and photothermal units. Chill portions (Fishman et al. 1987a, 1987b) were calculated using chillR (Luedeling et al. 2023). Accumulation of chill portions provides an index of progress towards fulfillment of potential vernalization requirements. Photothermal units were calculated according to Burghardt et al. (2015) such that each germination timing cohort accumulated units on the basis of hourly temperatures and day lengths with a base temperature rate of 4°C. We also determined the day length for each day of the study for the location of the screenhouse using chillR (Luedeling et al. 2023).

### Data collection

We measured phenological and morphological traits for all transplanted individuals. To quantify phenology, we censused individuals three times a week and recorded dates of first bud, first flower, and first fruit. The dates of these phenological stages were highly correlated so only first bud dates are considered in analyses here. We also recorded the height of each individual at these phenological stages. From these traits, we calculated time to first bud and first bud date for each individual. Time to first bud was calculated as the number of days between transplant date and date of first bud, an index of developmental speed. First bud date was calculated as the number of days from September 1 until first bud to standardize among germination timing cohorts. This metric represents the seasonal timing of first bud production.

To assess effects of germination timing on individual fitness, we recorded whether each plant flowered, and scored seed number and total seed mass for all flowering individuals. We then calculated first year fitness for each individual as the probability of flowering multiplied by the number of seeds produced. This metric is equivalent to lifetime fitness in annual species, but it does not account for reproduction in later years for biennials or perennials.

### Data analysis

#### Does flowering phenology respond to germination timing?

To examine phenological responses to germination timing, we fitted linear models to time to first bud, first bud date, and size at first bud. Each of these models included germination timing (transplant date coded as continuous), species, germination timing x species interaction, transplant height, bench, and population as predictors. Analysis of deviance tables were generated for fitted models to evaluate the significance of main effects (Table S4). Marginal means of the relationship of time to first bud, first bud date, and size at first bud with germination timing were then estimated for each species using the emmeans package (Lenth 2023).

#### Does germination timing affect fitness, either directly or indirectly through flowering phenology?

As with the phenology models, all fitness response models included germination timing (transplant date coded as continuous), species, their interaction, transplant height, bench, and population as predictors. To explore effects of germination timing on flowering probability, we fitted a generalized linear model with binomial error and a logit link function. Negative binomial generalized linear models (MASS package; Venables and Ripley 2002) were fitted to evaluate the relationships of number of seeds (for individuals that flowered) and first year fitness with germination timing whereas a linear model was fitted to model the same relationship for total seed mass. Analysis of deviance tables were generated for fitted models to evaluate the significance of main effects (Table S5) and then marginal means of the relationships between each fitness response and germination timing were estimated for each species using the emmeans package (Lenth 2023).

To decompose the effects of germination timing on fitness into direct effects and indirect effects through flowering phenology, we used structural equation models (SEMs) for time to first bud. Models were built separately for each species using piecewiseSEM (Lefcheck 2016). Prior analyses did not find significant differences among populations within species so populations were pooled for SEM analysis. The models consisted of (1) a linear model fit to the phenology response with germination timing (transplant date coded as continuous) and transplant height as predictors and (2) a negative binomial generalized linear model fit to number of seeds with the phenology response, germination timing, and transplant height as predictors. Tests of directed separation and Fisher’s C statistic were used to evaluate goodness of fit of the SEMs (Lefcheck 2016; Lefcheck 2021). The latent-theoretical approach was used to calculate standardized path coefficients for the linear phenology sub-model while the observation-empirical approach was used for the negative binomial fitness sub-model (Grace et al. 2018; Lefcheck 2021). Diagrams of the SEMs were generated using DiagrammeR (Iannone 2023).

#### How have responses of flowering time and fitness to germination timing diversified across the clade, and do patterns reflect climate of origin?

We evaluated how relationships between phenology, fitness, and germination timing were distributed across the phylogeny of these species to understand how these responses have diversified. The phylogenetic hypothesis for the Streptanthoid complex was previously generated by Cacho et al. (2014) using six single copy nuclear genes and two chloroplast regions based on Bayesian MCMC analyses that used three 50-million-generation independent runs with sampling every 5000 generations. We tested for phylogenetic signal in the marginal means estimated for each species from relationships between time to first bud, first bud date, probability of flowering, number of seeds, total seed mass, first year fitness and germination timing. Blomberg’s K was estimated using phytools (Revell 2012) with standard errors of the marginal means included in the estimations. Values of Blomberg’s K equal to one indicate that species variation in these relationships are indistinguishable from Brownian motion along the phylogeny (Blomberg et al. 2003). When values of K are greater than or less than one, there is more or less variation, respectively, among species in the relationships than expected given Brownian motion (Blomberg et al. 2003).

We tested whether slopes of relationships between time to first bud, probability of flowering, number of seeds, first year fitness, and germination timing were related to species’ climate using phylogenetic generalized linear models (Orme et al. 2023). Six climate variables were extracted from the California Basic Characterization Model (Flint and Flint 2014) for species’ locations across their ranges taken from herbarium records from the Consortium of California Herbaria (CCH2 Portal 2023; Table S6). Average annual values of climate water deficit, precipitation, minimum and maximum temperatures, as well as inter-annual variability in precipitation (coefficient of variation) and temperature (standard deviation) over 25 years (1991-2015) were calculated and a principal component analysis was used to reduce dimensionality of the data (Table S6). The first principal component was used in the models as it was strongly associated with all climate variables except variation in minimum and maximum temperatures. This component contrasts cool, wet climates with less variation in precipitation (high values of PC1) from warm, dry climates with more variable precipitation (Table S6).

We also used phylogenetic generalized linear models to test whether species with more specialized germination niches, those germinating in a narrower seasonal window with respect to rainfall onset, would also be more specialized in their post-germination niche requirements. If so, germination specialists should exhibit greater fitness reductions when forced to germinate outside their usual seasonal germination niche, than species that germinate across a broader range of rainfall onset dates. In a previous study, we showed that germination fraction declined with later rainfall onset date (Worthy et al. 2023). Here, we used the slope of this relationship as an index of germination specialization for each species; steeper slopes indicated narrower germination niches concentrated earlier in the season. To test our prediction, we asked whether such germination specialization was associated with species declines in fitness, number of seeds produced or first year fitness, with later germination timing in the present study. For this analysis, slopes of germination fraction for the species CAIN and CAAN, estimated in Worthy et al. (2023), were averaged over two populations.

Lastly, we used phylogenetic generalized linear models to test the hypothesis that species with greater acceleration of flowering in later cohorts would better maintain fitness across a range of germination dates, as expected if this compensatory plasticity is adaptive. We tested this hypothesis by evaluating relationships between slopes of time to first bud against germination timing and slopes of three fitness metrics, flowering probability, number of seeds produced, and first year fitness against germination timing.

## Results

### Does flowering phenology respond to germination timing?

Later germinating cohorts encountered different seasonal environments, including decreases in the amount of chilling and photothermal units individuals accumulated during the growing season and variation in day length (Figure S2). Effects on reproductive phenology of these shifts in seasonal conditions with germination timing varied among species. In all species, the number of days to production of the first bud decreased significantly with later germination timing (Figure 1; Table S7). Nevertheless, in all species except STTO which did not flower in later cohorts, individuals that germinated in later rain events produced their first bud later in the season than individuals that germinated in earlier rain events (Figure S3; Table S8). Thus, acceleration of bud production in later germination cohorts partially synchronized reproductive timing with earlier cohorts but did not completely prevent reproductive delays. Species differed in the degree of this compensatory plasticity of days to bud to germination timing (Figure 1), and consequently in the degree of synchronization of budding date across germination cohorts (Figure S3). Species with greater acceleration of budding in later season germination cohorts exhibited more synchronization of budding date with respect to germination timing (Figures 1; S3). In four species, plants from later germination cohorts were significantly shorter when producing their first bud than individuals in earlier cohorts (Figure S4; Table S9).

**Figure 1.**
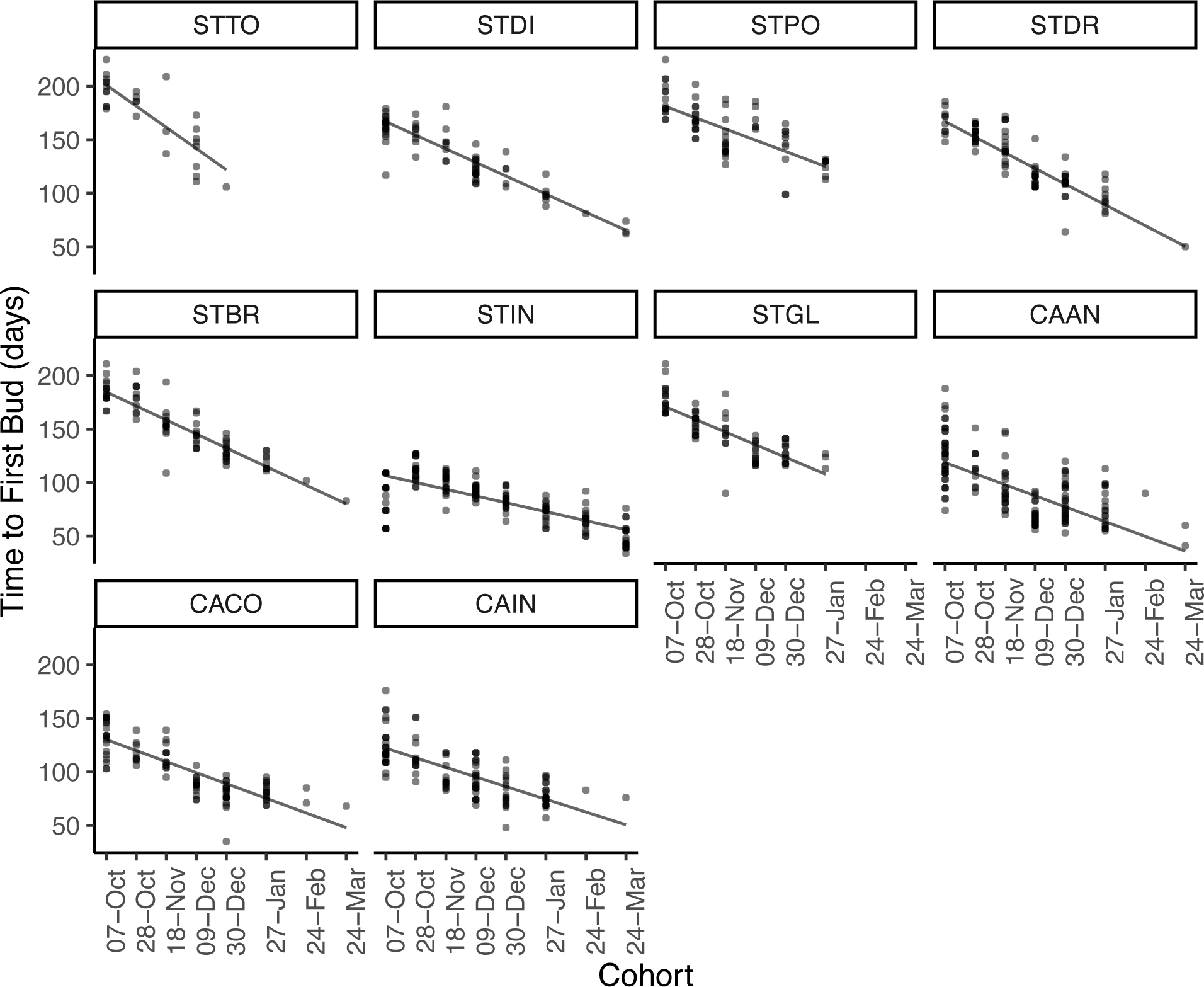
Relationships between amount of time to first bud (days) and germination timing. Time to first bud was calculated as the number of days between individual transplant date and date of first bud. Regression lines represent significant, negative relationships where individuals that germinated in later cohorts took less time to produce their first bud. Points represent observed time to first bud for individuals of each species in each cohort. Panels are ordered by species’ phylogenetic relationships.

### Does germination timing affect fitness, either directly or indirectly through flowering phenology?

Germinating later in the season translated, for many species, into reduced fitness. For five species, the probability of flowering significantly decreased as the timing of germination came later in the season (Figure 2; Table S10). The facultative biennial STTO failed to flower entirely in the last four germination cohorts, where chilling accumulation was likely insufficient to satisfy previously reported requirements for vernalization (Gremer et al. 2020a) (Figure 2; S5). Four other species also showed significant declines in probability of flowering in later cohorts, plausibly again due to insufficient chilling. One species, STDR, showed a significant increase in the probability of flowering with later germination, potentially due to high pre-reproductive mortality of individuals in the first cohort that resulted in low sample sizes for early cohort estimates (Figure S5). Three species showed no significant effect of germination timing on flowering probability (Figure 2).

**Figure 2.**
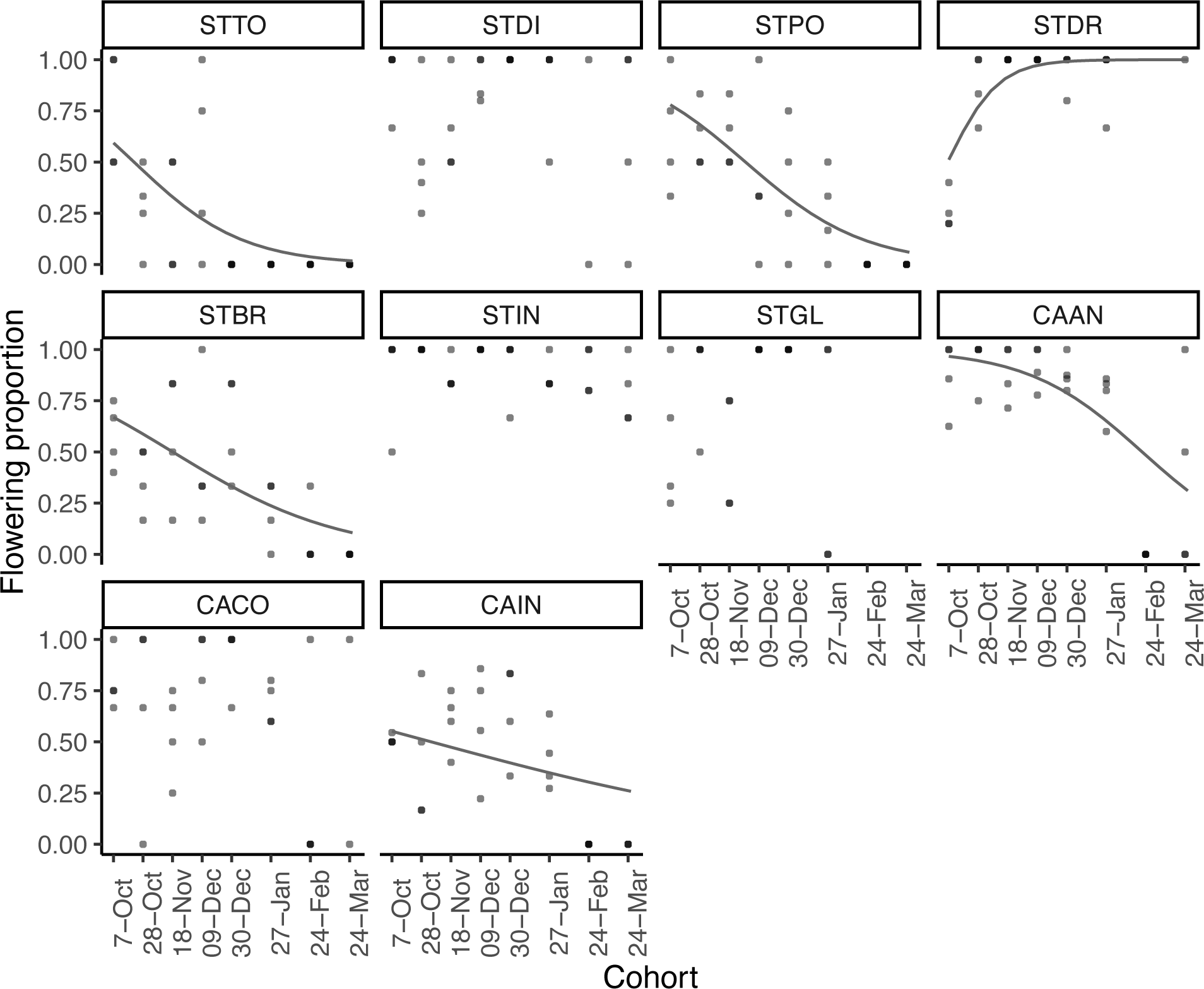
Relationships between probability of flowering and germination timing. Regression lines represent significant relationships (back transformed from logit scale) where later germinating individuals had lower probability of flowering, except for STDR which had high pre-reproductive mortality of individuals in the first cohort (Figure S5). Points represent mean proportions of flowering individuals of each species in each cohort on each of four benches. Panels are ordered by species’ phylogenetic relationships.

Flowering individuals of six species produced significantly fewer seeds when germinating later in the season (Figure 3; Table S11) and four of these species also had significantly lower total seed mass with later germination timing (Figure S6; Table S12). Overall, seven of the ten species showed significant decreases in their first year fitness as the timing of rainfall events and subsequent germination occurred later in the season (Figure S7; Table S13).

**Figure 3.**
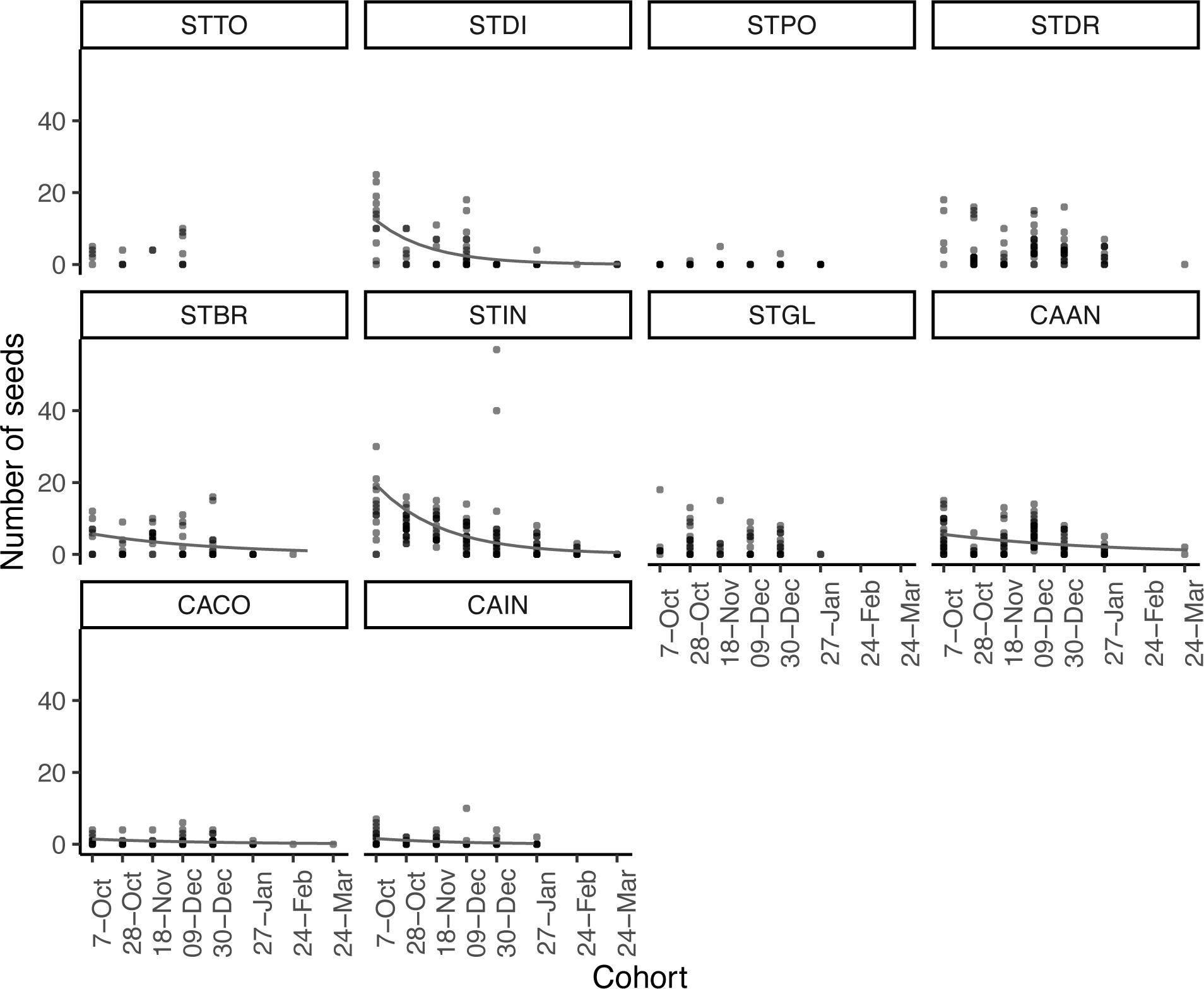
Relationships between number of seeds produced and germination timing. Regression lines represent significant, negative relationships where later germinating individuals produced fewer numbers of seeds. Points represent the number of seeds for individuals of each species in each germination timing cohort. Many individuals included in this analysis flowered but did not produce any seeds (n = 389). Panels are ordered by species’ phylogenetic relationships.

Structural equation models revealed that germination timing influenced seed production both directly and indirectly through its effects on flowering phenology (Figure 4; Table S14). Later germination timing had a significant, direct, negative effect on seed number for all species except for STPO, a nickel-hyperaccumulator (Reeves et al. 1981), which had lower overall seed set throughout the experiment (Figure 4). This negative effect was partially counteracted by a positive indirect effect through time to first bud, suggestive of adaptive phenotypic plasticity (Table S14): later germination resulted in significantly fewer days to bud, which in turn had a significant positive effect on seed number in all species except STPO and STIN (Figure 4). Thus, acceleration of flowering in later cohorts partially ameliorated the fitness impacts of later germination. However, the relative magnitude of direct and indirect effects, and thus the total effect of germination timing on fitness, varied among species (Figure 4; Table S14).

**Figure 4.**
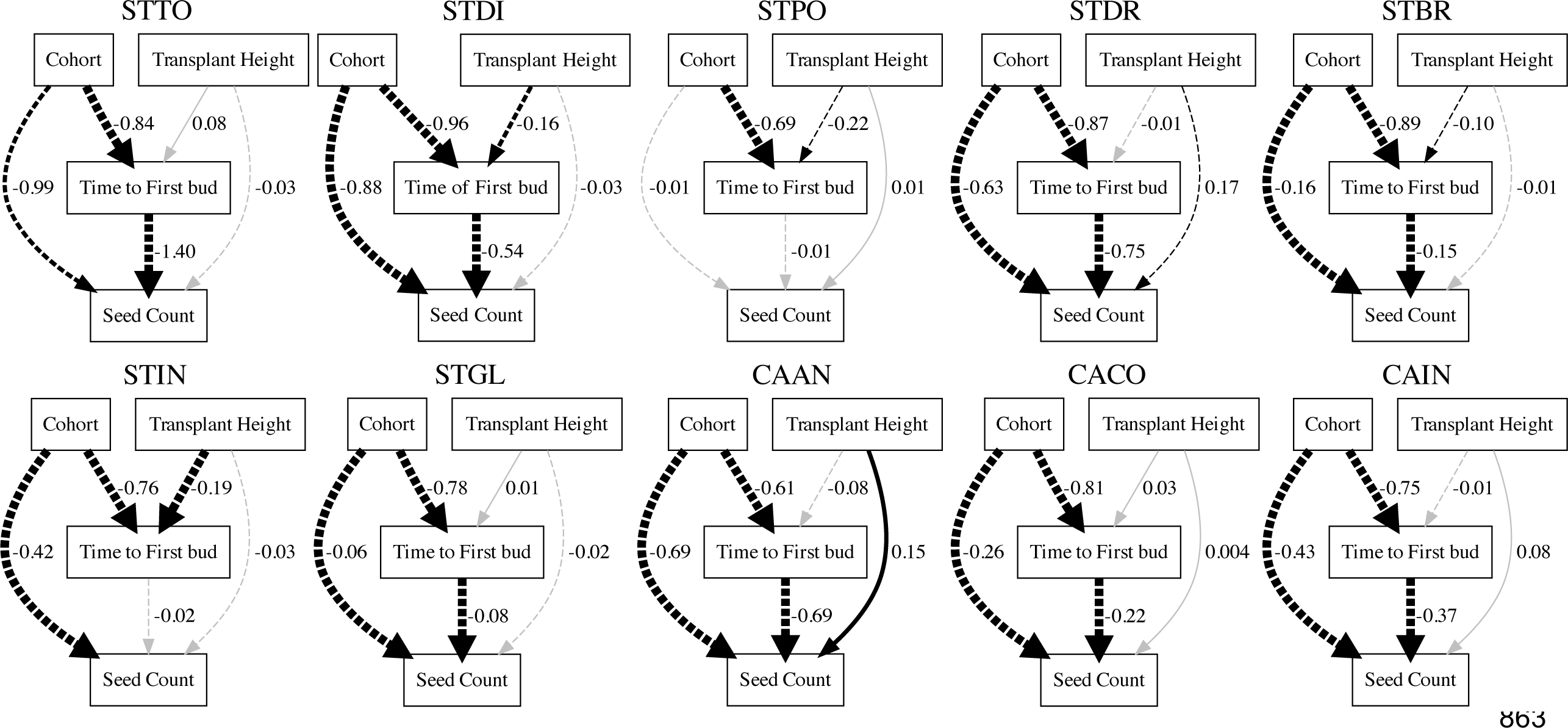
Results of structural equation models testing for direct effects of germination timing (cohort) on fitness (seed count) and indirect effects through phenology, time to first bud (days). Time to first bud was calculated as the number of days between individual transplant date and date of first bud. Numbers are standardized partial regression coefficients, and the width of the arrows is scaled based on their level of significance. Solid and broken lines represent positive and negative relationships, respectively. Line color represents significant (black) and non-significant (gray) relationships. Species are in order of phylogenetic relationships.

### How have responses of flowering time and fitness to germination timing diversified across the clade, and do patterns reflect climate of origin?

Species’ reproductive phenology and fitness responses to the timing of germination varied substantially across the clade. We did not find significant phylogenetic signal for responses of time to first bud or first bud date (K = 1.18, p = 0.15; Figures 5; S8) to germination timing with values consistent with Brownian motion; note that these phenological traits are linear transformations of one another so the statistics are identical. Bloomberg’s K for responses of flowering probability (K = 0.97, p = 0.91; Figure S8) and number of seeds produced (K = 1.07, p = 0.84; Figure 5) to germination timing were consistent with Brownian motion. In contrast, Blomberg’s K values less than one for seed mass (K = 0.77, p = 0.39; Figure S8) and first year fitness (K = 0.68, p = 0.99; Figure S8) suggest that some fitness responses to germination timing may be more evolutionarily labile than phenological responses.

**Figure 5.**
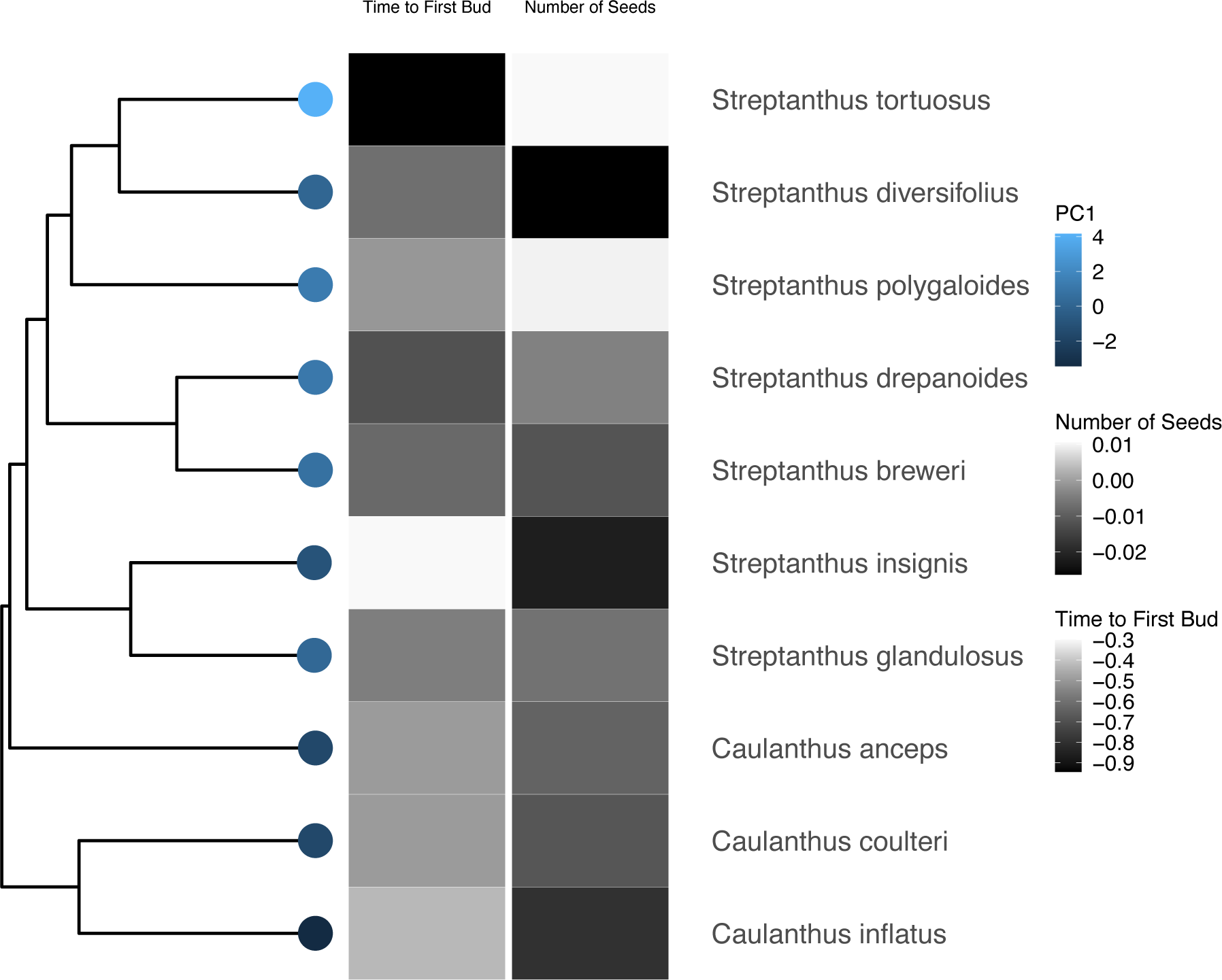
Slopes of relationships of time to first bud and number of seeds produced with timing of germination displayed across the phylogeny. Time to first bud was calculated as the number of days between individual transplant date and date of first bud. Phylogenetic signal of these relationships was evaluated with Blomberg’s K: time to first bud (K = 1.18, p = 0.15); number of seeds (K = 1.07, p = 0.84). Scores associated with the first principal component of an analysis of average yearly climate for 1991-2015 for species’ locations is displayed (Table S6).

After accounting for phylogenetic relatedness, the responses of time to first bud and number of seeds to germination timing were significantly related to the first principal component of species’ range-wide 25-year average annual climatic conditions (Figure 6A, 6C, Table S15), suggesting a role of climate adaptation in species divergence. Species with steeper slopes of time to first bud (i.e. those whose reproductive phenology was more plastic to germination timing) were associated with lower climate water deficient (CWD), lower variability in precipitation, and lower maximum temperature, and higher mean annual precipitation (Figure 6A, Table S6). These species also had shallower decreases in number of seeds produced with later germination timing (less negative slopes; Figure 6C). Slopes of flowering probability and first year fitness against germination time were unrelated to climate (Figure 6B, 6D, Table S15).

**Figure 6.**
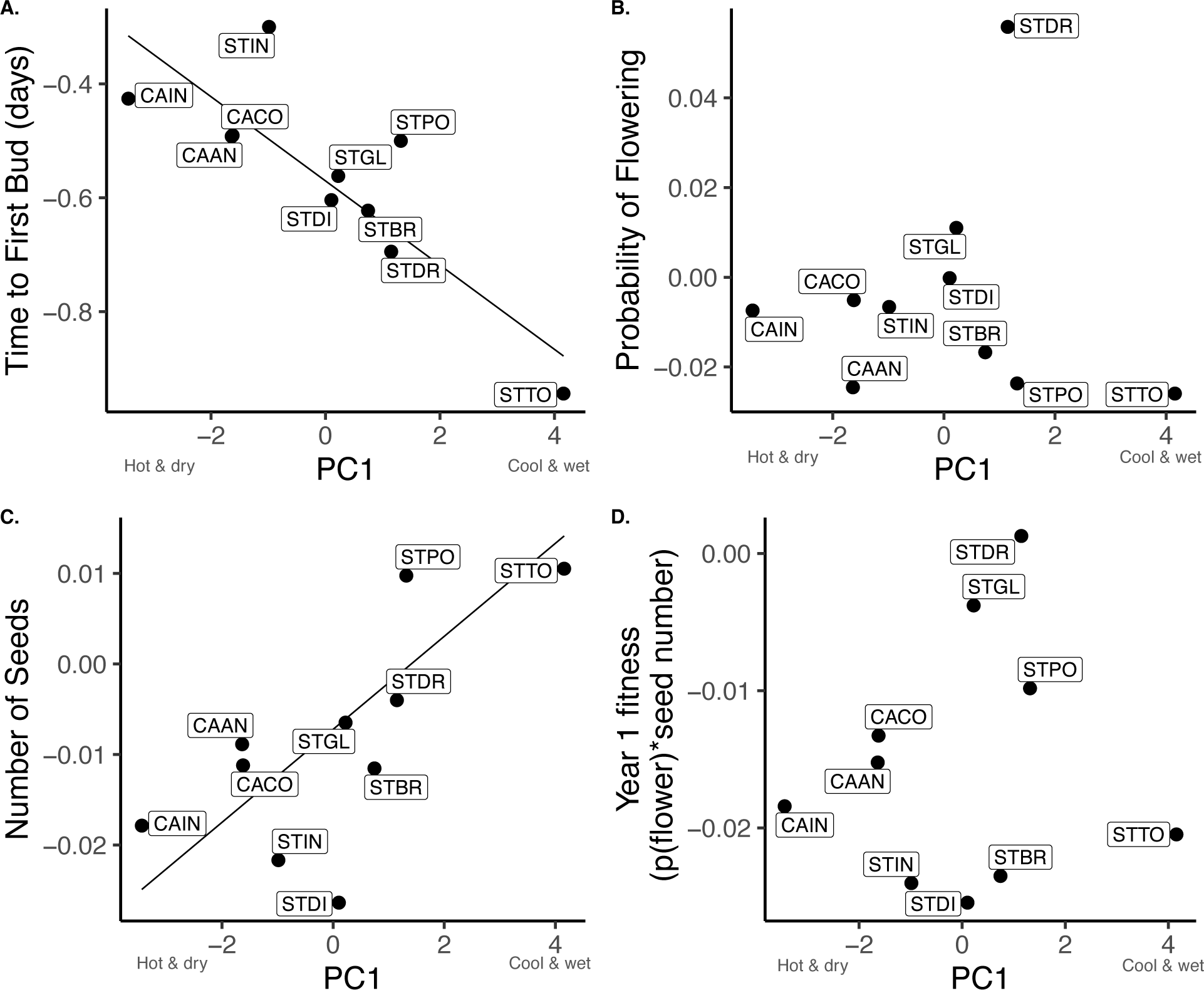
Relationships between phenology and fitness responses to germination timing and 25-year average annual climate variables tested using phylogenetic generalized linear models. Separate models were built for slopes of relationships between **(A)** time to first bud, **(B)** probability of flowering, **(C)** number of seeds produced, **(D)** first year fitness and germination timing. Principal component analysis was used to reduce dimensionality of the climate variables (Table S6). Most climate variables were associated with PC1, so it was used as the predictor in these models. Points represent slope estimates for each species.

We had hypothesized that species with narrower seasonal germination niches observed in our previous study (Worthy et al. 2023) might have greater fitness sensitivity when forced to experience seasonal conditions under which they normally would avoid germinating. However, results of phylogenetic generalized linear models did not support that hypothesis: steeper declines in germination proportion in later seasonal cohorts (i.e. narrower seasonal germination niches) were not significantly associated with steeper declines in fitness with later germination (narrower post-germination seasonal niches; Figure S9; Table S16) across species.

We also tested the hypothesis that compensatory plasticity of days to bud (or stasis of bud date) to germination timing would be associated with stabilization of fitness across germination cohorts. Accounting for phylogenetic relatedness, we observed a marginally significant negative relationship between species’ phenological plasticity of days to bud (e.g. the slope of the relationship between time to first bud and germination timing) and response of seed number to germination timing (Figure S10A; Table S17). Species with steeper decreases in time to first bud - and consequently smaller delays in bud date - had shallower declines in seed number with later germination timing, as predicted if acceleration of flowering in later cohorts is adaptive, compensatory plasticity. However, the relationships between phenological plasticity, flowering probability (Figure S10B), and first year fitness were not significant (Figure S10C).

## Discussion

The seasonal timing of seed germination is a niche construction trait that determines the environmental conditions that plants experience during the growing season, thus shaping phenology and fitness (Donohue 2005; Donohue et al. 2005). As climate change alters seasonal environments, the resulting changes in seasonal germination timing may critically influence plant population performance (Kimball et al. 2011, Levine et al. 2011; Gremer et al. 2020b, Martínez-Berdeja et al. 2023). Our results reveal that later germination due to later onset of the rainy season in Mediterranean climates has significant impacts on flowering phenology, fitness, and potential resilience of plant populations to climate change, but these impacts vary among species across the *Streptanthus/Caulanthus* clade. A critical component of these differences is species’ variation in the degree of compensatory phenological plasticity: the ability of later germinating plants to accelerate flowering, thus reducing delays in flowering date and stabilizing fitness responses to germination timing. This compensatory plasticity has diversified across the clade with a signal of climate adaptation. Fitness responses to germination timing, in contrast, appear more evolutionarily labile, with limited evidence of climate adaptation. As climate change brings later average onset of the California rainy season (Luković et al. 2021), compensatory phenological plasticity may partially mitigate fitness impacts of delayed germination, but the majority of species are likely to suffer decreases in population fitness.

### Does flowering phenology respond to germination timing?

Germination timing often has cascading effects on flowering phenology, which may be mediated by availability of and species’ responsiveness to seasonal cues (Donohue et al. 2002, 2010; Wilczek et al. 2009; Burghardt et al. 2015; Miryeganeh et al. 2018; Gremer et al. 2020a; Olliff-Yang-and Ackerly 2021). In our study, later germination cohorts generally flowered in fewer days, but species differed in the degree of compensatory plasticity; species with greater plastic acceleration of days to bud in response to later germination had more synchronized flowering dates across germination cohorts. However, compensatory plasticity of days to bud was insufficient to synchronize flowering date completely. Similarly, plastic acceleration of flowering with later germination does not entirely synchronize flowering date in *Arabidopsis thaliana* (Wilczek et al. 2009, Miryeganeh et al. 2018, Martinez-Berdeja et al. 2023, Huang et al. 2024).

Since all species experienced identical seasonal environments in our experiment, variation in phenological responses indicates that species differed in responses to seasonal cues. Day length changed dramatically throughout the study while accumulation of chilling hours declined drastically in later germination cohorts, accompanied by significant declines in the probability of flowering in five species. One of these species was the facultative biennial STTO, in which we have experimentally demonstrated a vernalization requirement for flowering (Gremer et al. 2020a). That finding suggests that failure to flower in late germinating cohorts of the other related species may also stem from unsatisfied vernalization requirements. However, the adaptive value of this requirement for fall-germinating winter annuals is unclear. Facultative vernalization requirements can prevent premature flowering in unfavorable winter conditions and synchronize flowering in spring, but flowering eventually occurs in the absence of chilling (Wilczek et al. 2009, Miryeganeh et al. 2018, Huang et al. 2024). However, failure to flower in spring due to insufficient vernalization means zero fitness for winter annuals, and spring germination may then be a lethal strategy. These temperature cues may interact with other factors, such as photoperiod and plant ontogeny or size, to influence flowering phenology (Donohue et al. 2010; Burghardt et al. 2015). Although our experimental design cannot distinguish the relative contributions of seasonal cues to species response differences, the results provide hypotheses to inform future experiments manipulating specific cues in controlled environments (e.g. Friedman and Willis 2013, Burghardt et al. 2016, Wolkovich et al. 2022).

### Does germination timing affect fitness, either directly or indirectly through flowering phenology?

Germination timing is often under strong selection (Kalisz 1986; Donohue et al. 2005; Verdú and Traveset 2005; Picó 2012; Postma and Agren 2016, 2022; Zachello et al. 2020; Martínez-Berdeja et al. 2023). For annual plants in seasonal environments, late germination may affect fitness in two ways: through exposure to less favorable conditions at the time of emergence, and by decreasing the window of time available for growth and reproduction before the end of the growing season. Indeed, for all species in our experiment, these combined direct effects of later germination reduced seed production. The negative fitness impact of late germination was, however, partially mitigated by positive indirect effects of germination timing through its accelerating effect on phenology. Thus, acceleration of reproduction in later germination cohorts seems to be a form of adaptive plasticity that compensates for the negative direct effects of later germination. The degree of compensation and the balance of direct and indirect effects varied among species, resulting in the observed differences in fitness responses to germination timing. Direct and indirect effects of seasonal timing have also been observed in terrestrial and aquatic migratory species where changes to environmental conditions, such as warming temperatures, have altered the timing of migration with direct effects on fitness and indirect effects through, for example, first egg date in birds (Both and Visser 2001; Winkler et al. 2014; Inouye 2022). It is important to note that our fitness metrics did not account for effects of seasonal selective factors, such as pollinator or mate availability or herbivore abundance, that may be important in the field (Miryeganeh et al. 2018; Kehrberger and Holzschuh 2019, Kudo and Cooper 2019).

### How have responses of flowering time and fitness to germination timing diversified across the clade, and do patterns reflect climate of origin?

Phylogenetic analyses suggest a signature of climate adaptation in the evolutionary diversification of phenological plasticity and seed production responses to rainfall onset across the Streptanthoid clade. Species from drier, warmer, more variable habitats in southern California accelerated flowering less in later cohorts and had less synchronized flowering dates than northern species. This finding suggests that compensatory phenological plasticity to germination timing may have evolved as the clade diversified out of the desert into cooler, wetter, more stable environments, with longer growing seasons. The desert species flowered rapidly even in the earliest germination cohorts, suggesting a constant developmental program across seasonal environments. This risk-averse, constitutively early flowering strategy may be adaptive in the short growing season and unpredictable rainfall of their home climates (Wesselingh et al. 1997; Metcalf et al. 2003; Rees et al. 2004; Austen et al. 2017). In contrast, Streptanthoid species from cool, wet environments were more plastic in flowering phenology (Pearse et al. 2020). Early germination cohorts flowered at a later age and larger size, which may help to maximize fitness in a longer growing season with early rainfall. Later cohorts accelerated development to flower at an earlier age and smaller size, synchronizing flowering date and allowing reproduction despite a shorter growing season. Responses of other fitness traits, first year fitness and seed mass, to germination timing were more evolutionarily labile, with no climate association and no correlation with phenological plasticity across the clade. Fitness and life history metrics, in general, have previously been noted as less evolutionarily conserved (Burns et al. 2010; Salguero-Gómez et al. 2016; Che-Castaldo et al. 2018), connected to colonization of new habitats (Herben et al. 2014) and higher intraspecific variation (Healy et al. 2019).

Consistent with our finding that compensatory phenological plasticity mitigated the direct negative fitness effects of later germination within most of our study species, we found that species with greater acceleration of days to bud in later cohorts had shallower declines in seed number after correcting for phylogeny. Such plasticity may buffer fitness declines in the face of delayed rainfall onset. Indeed, a phylogenetic analysis by Willis et al. (2008) showed that species with greater plasticity of flowering time to spring temperatures decreased less in abundance over time. Another strategy to mitigate the effects of shifting germination timing would be to avoid germinating outside favorable environmental windows. This strategy would require specialization on a narrower range of germination conditions, with higher fitness costs expected for species with this strategy forced to germinate in unfamiliar seasonal environments. We did not find support for this strategy here; responses of germination and post-germination fitness components were uncorrelated, suggesting that they have evolved independently across the phylogeny. Thus, compensatory phenological plasticity is likely the primary mechanism to buffer effects of variation in germination timing in this system.

### Can variation among species in responses to germination timing lead to differential vulnerability to climate change?

Whether or not phenotypic plasticity can buffer fitness across changing environments is an important question for predicting population resilience to climate change (Ghalambor et al. 2007; Chevin and Lande 2010; Chevin et al. 2010; Nicotra et al. 2010; Vedder et al. 2013; Valladares et al. 2014; Anderson and Gezon 2015; Duputié et al. 2015; Kingsolver and Buckley 2017; DeMarche et al. 2018; Scheiner et al. 2020; Gauzere et al. 2020). Numerous studies have focused on plastic phenological responses to changing spring environments, such as early budburst or flowering in response to warmer springs and earlier snowmelt (Willis et al. 2008; Anderson et al. 2012; Anderson and Gezon 2015; Wadgymar et al. 2018; Zettlemoyer et al. 2024). Less attention has been paid to plant responses to changing fall environments (but see Kimball et al. 2010; Martinez-Berdeja et al. 2023; Worthy et al. 2023). In desert and Mediterranean climates, later onset of fall rains (Luković et al. 2021) means that seeds are germinating in cooler conditions (Kimball et al. 2010; Worthy et al. 2023) and may have less time to complete reproduction before the end of the growing season. This shorter growing season may therefore select for a faster transition to flowering, but at the expense of smaller size at reproduction, and consequent lower fecundity (Cohen 1976).

In our study, species differences in degree of phenological compensation resulted in different sensitivities of seed production to seasonal germination delays; over half of the species significantly declined in fecundity with later germination. For these species, compensatory phenological plasticity partially mitigated fitness costs, but was insufficient to stabilize fitness in the face of delayed germination. Thus, acceleration of flowering may be an adaptive response in short seasons but may still reduce population mean fitness (Colautti et al. 2017). Plasticity of flowering time to germination timing may allow plants to maximize fecundity in years with early rainfall onset while mitigating fitness costs of shorter growing seasons in years with late rainfall, buffering mean population fitness across years. However, as the mean rainfall date moves later in the year, mean population fitness will decline. This finding adds to the growing body of evidence that adaptive phenological plasticity may be insufficient to fully maintain population fitness in the face of climate change (Franks et al. 2014; Duputié et al. 2015; Colautti et al 2017; Zettlemoyer et al. 2024).

Together, these findings suggest that populations of many Streptanthoid species are vulnerable to future delays in rainfall onset with climate change. Species with vernalization requirements for flowering are especially likely to suffer greater impacts of later rainfall due to reproductive failure, especially if future warming temperatures reduce the seasonal accumulation of chilling hours (Cook et al. 2012; Ettinger et al. 2020; Anderson 2023). For such species, a formerly adaptive cue of seasonal conditions for flowering becomes maladaptive under novel climate conditions, and selection may act to weaken vernalization requirements in the future. We have previously demonstrated that germination declines with later rainfall onset, although species differ across the clade in the magnitude of this decline (Worthy et al. 2023). This finding, combined with our current results, suggests that the impact of delayed rainfall onset on fitness will have cascading effects across life stages. Taken together, these findings suggest that species resilience to climate change across the Streptanthoid clade may depend upon the cascading fitness consequences of phenological responses to seasonal cues across life stages.

## Acknowledgments

We thank Chloe Abbasi, Eda Ceviker, Bryce Johnson, Hugo Mahatdejkul, Shannon Reilly, Emily Rushka, Samantha Swan, and Yasmiin Wadhwani for assistance with experimental setup, maintenance, and data collection. This research was supported by funding from the National Science Foundation grant DEB-1831913 awarded to J.R. Gremer, J. Schmitt, S.Y. Strauss, and J. N. Maloof, a USDA Hatch Grant CA-D-PLB-2795-H awarded to J. N. Maloof., a USDA Hatch grant CA-D-EVE-3515-H-1002936 awarded to J. Schmitt, and U.C. Davis.

## Author Contributions

J.R.G, J.S., S.Y.S., and J.N.M conceived and designed the study. S.J.W., A.M, and S.R.A performed the experiment and gathered the data. S.J.W., J.R.G., S.Y.S., and J.S designed the data analyses. S.J.W. and J.R.G. analyzed the data with assistance from J.S., S.Y.S, and S.R.A. S.J.W., S.R.A., and J.S. wrote the manuscript with contributions from all other authors. All authors contributed to development of ideas, analyses, and interpretation of results.

## Data and Code Availability

All data and code necessary to reproduce the results in this article are available at https://github.com/StreptanthusDimensions/Germination.Fitness. A Zenodo doi will be obtained for this material upon manuscript acceptance.

