## Supplementary Material for "Accelerated phenology fails to buffer fitness loss from delayed rain onset in a clade of wildflowers"

**Table S1.** Species (10) and populations (12) included in this study from the *Streptanthus* and *Caulanthus* genera of Brassicaceae along with their abbreviation used in figures in the manuscript. General population location and seed source information are also given. A screenhouse has a clear plastic roof, but screened sides so that watering can be controlled, but plants experience seasonal temperature fluctuations and changes in ambient light. Average annual precipitation (mm) over 30 years (1991-2020) was used to group species into watering regimes. Data downloaded from PRISM (PRISM Climate Group 2014). Sample size ranges across all cohorts combined are presented for each population.

| Species | Abbreviation | Population | Seed Source | Precipitation | Sample Size Range |
|---|---|---|---|---|---|
| *Caulanthus anceps* | CAAN1 | Carrizo Plain | Screenhouse | 225 | 4 - 24 |
| *Caulanthus anceps* | CAAN2 | Ballinger Canyon | Field | 309 | 1 - 24 |
| *Caulanthus coulteri* | CACO | Frazer Road | Screenhouse | 341 | 2 - 24 |
| *Caulanthus inflatus* | CAIN3 | Ballinger Canyon | Field | 309 | 3 - 24 |
| *Caulanthus inflatus* | CAIN4 | Hurricane Road | Field | 230 | 1 - 24 |
| *Streptanthus breweri* | STBR | Turtle Rock | Screenhouse | 803 | 12 - 24 |
| *Streptanthus diversifolius* | STDI | Big Sandy | Screenhouse | 758 | 2 - 24 |
| *Streptanthus drepanoides* | STDR | Lime Saddle | Screenhouse | 1195 | 0 - 24 |
| *Streptanthus glandulosus* | STGL | Bartlett Springs | Field | 1171 | 0 - 24 |
| *Streptanthus insignis* | STIN | Panoche Road | Screenhouse | 288 | 21 - 24 |
| *Streptanthus polygaloides* | STPO | Magalia | Field | 1637 | 11- 24 |
| *Streptanthus tortuosus* | STTO | Table Mountain 2 | Field | 1066 | 5 - 17 |

**Table S2.** Temperature regime experienced by each germination timing cohort in growth chambers (E7/2 Conviron, Winnipeg, Manitoba, Canada) with a 12-h daylight cycle for 17 days. Temperatures were adjusted to increase germination.

| **Germination Timing Cohort** | **Temperature Conditions** |
|---|---|
| Cohort 1: All species | 25°C for first 9 days and 22°C for remaining 8 days |
| Cohort 2: All species | 22°C for all days |
| Cohort 3 – Cohort 8: CAAN, CACO, CAIN, STBR, STDI, STIN,STPO, STTO | 25°C for all days |
| Cohort 3 – Cohort 8: STDR and STGL | 25°C during the day and 15°C at night for all days |

**Table S3.** Watering regime species experienced during the study to replicate variation among species in field precipitation amounts. Previous experiments had shown that species from drier climates got fungal disease if watered too much. Watering groups were determined using species' 30-year average annual precipitation amounts (Table S1). When necessary due to excess heat, additional water was supplied in 30-sec boosts, with proportional increases across the watering groups.

| Watering Group | Total Watering Time per Week (min.) | Total Water per Week (mL) |
|---|---|---|
| Low (225-340 mm) | 3 | 49.50 |
| Medium (630-803 mm) | 3.5 | 57.75 |
| High (1065-1637 mm) | 4.5 | 75.25 |

**Table S4.** Analysis of deviance tables generated after fitting models of three phenology metrics. Basic linear models were used to model first bud date, time to first bud, and size at first bud (height). All three of these models included germination timing (transplant date as continuous), species, their interaction, transplant height, bench, and population as predictors. For each model, the degrees of freedom (Df), sum of squares (Sum Sq), mean squares (Mean Sq), F statistic (F-value), and P-value are reported. P-values in bold were significant ($p < 0.05$).

| | Df | Sum Sq | Mean Sq | F-value | P-value |
|---|---|---|---|---|---|
| First Bud Date | | | | | |
| Germination timing | 1 | 366681 | 366681 | 1442.62 | **< 2.2e-16** |
| Species | 9 | 582520 | 64724 | 254.64 | **< 2.2e-16** |
| Transplant height | 1 | 2129 | 2129 | 8.37 | **0.003889** |
| Bench | 3 | 1822 | 607 | 2.39 | 0.067444 |
| Population | 2 | 173 | 87 | 0.34 | 0.711099 |
| Germination timing*Species | 9 | 33577 | 3731 | 14.68 | **< 2.2e-16** |
| Time to First Bud | | | | | |
| Germination timing | 1 | 610528 | 610528 | 2401.98 | **< 2.2e-16** |
| Species | 9 | 582520 | 64724 | 254.64 | **< 2.2e-16** |
| Transplant height | 1 | 2129 | 2129 | 8.37 | **0.003889** |
| Bench | 3 | 1822 | 607 | 2.39 | 0.067444 |
| Population | 2 | 173 | 87 | 0.34 | 0.711099 |
| Germination timing*Species | 9 | 33577 | 3731 | 14.68 | **< 2.2e-16** |
| Size at First Bud | | | | | |
| Germination timing | 1 | 138.4 | 1384.25 | 193.11 | **< 2.2e-16** |
| Species | 9 | 5275.0 | 586.11 | 81.76 | **< 2.2e-16** |
| Transplant height | 1 | 382.4 | 382.36 | 53.34 | **8.37e-13** |

| | | | | | |
|---|---|---|---|---|---|
| Bench | 3 | 21.9 | 7.31 | 1.02 | 0.3835 |
| Population | 2 | 12.0 | 6.00 | 0.84 | 0.4335 |
| Germination timing*Species | 9 | 871.9 | 96.88 | 13.52 | **< 2.2e-16** |

**Table S5.** Analysis of deviance tables generated after fitting models of four fitness responses to the timing of germination. First year fitness was calculated as the probability of flowering multiplied by the number of seeds produced. A generalized linear model was used to model flowering probability. Negative binomial generalized linear models were used to model the number of seeds and first year fitness while a basic linear model was fit to total seed mass. All four of these models included germination timing (transplant date as continuous), species, their interaction, transplant height, bench, and population as predictors. For each model, the degrees of freedom (Df), sum of squares (Sum Sq), mean squares (Mean Sq), F statistic (F-value), and P-value are reported. P-values in bold were significant ($p < 0.05$).

| | Df | Sum Sq | Mean Sq | F-value | P-value |
|---|---|---|---|---|---|
| Flowering probability | | | | | |
| Germination timing | 1 | 65.18 | 1342 | 1712.10 | **6.838e-16** |
| Species | 9 | 282.67 | 1333 | 14.29.40 | **< 2.2e-16** |
| Transplant height | 1 | 18.25 | 1332 | 1411.20 | **1.936e-05** |
| Bench | 3 | 4.82 | 1329 | 1406.30 | 0.1852 |
| Population | 2 | 2.87 | 1327 | 1403.50 | 0.2387 |
| Germination timing*Species | 9 | 81.14 | 1318 | 1322.30 | **9.599e-14** |
| Number of Seeds | | | | | |
| Germination timing | 1 | 95.07 | 838 | 1121.74 | **< 2.2e-16** |
| Species | 9 | 247.84 | 829 | 873.91 | **< 2.2e-16** |
| Transplant height | 1 | 1.74 | 828 | 872.17 | 0.1871 |
| Bench | 3 | 0.97 | 825 | 871.20 | 0.8085 |
| Population | 2 | 1.81 | 823 | 869.39 | 0.4046 |
| Germination timing*Species | 9 | 34.41 | 814 | 834.98 | **7.572e-05** |
| Total Seed Mass | | | | | |

| | | | | | |
|---|---|---|---|---|---|
| Germination timing | 1 | 0.000347 | 0.000347 | 69.22 | **3.699e-16** |
| Species | 9 | 0.000803 | 0.000089 | 17.81 | **< 2.2e-16** |
| Transplant height | 1 | 0.000015 | 0.000015 | 3.05 | 0.08124 |
| Bench | 3 | 0.000004 | 0.000001 | 0.23 | 0.87292 |
| Population | 2 | 0.000001 | 0.000001 | 0.11 | 0.89407 |
| Germination timing*Species | 9 | 0.000264 | 0.000029 | 5.85 | **6.351e-08** |
| First Year Fitness | | | | | |
| Germination timing | 1 | 164.85 | 1341 | 1351.52 | **< 2.2e-16** |
| Species | 9 | 348.33 | 1332 | 1003.19 | **< 2.2e-16** |
| Transplant height | 1 | 9.72 | 1331 | 993.47 | **0.001827** |
| Bench | 3 | 2.85 | 1328 | 990.62 | 0.414878 |
| Population | 2 | 2.26 | 1326 | 988.36 | 0.322956 |
| Germination timing*Species | 9 | 26.29 | 1317 | 962.07 | **0.001830** |

**Table S6.** Principal component analysis of average yearly climate for 1991-2015 for species' locations taken from herbarium records from the Consortium of California Herbaria. Climate variables include climate water deficit (CWD), precipitation (PPT), minimum and maximum temperature (Tmin, Tmax, respectively), as well as inter-annual variability in precipitation (coefficient of variation, PPT_CV) and temperature (standard deviation of Tmin and Tmax, Tmin_SD and Tmax_SD, respectively). Data source was Flint and Flint, (2014). Cumulative proportion of variance explained by PC1 and PC2 is also presented.

| **Variable** | **PC1** | **PC2** |
|---|---|---|
| CWD | -0.47 | 0.03 |
| PPT | 0.46 | -0.04 |
| PPT_CV | -0.44 | 0.22 |
| Tmin | -0.35 | -0.17 |
| Tmin_SD | -0.14 | 0.68 |
| Tmax | -0.46 | -0.17 |
| Tmax_SD | 0.14 | 0.65 |
| Cumulative Proportion | 0.62 | 0.88 |

**Table S7.** Results from the linear model with time to first bud (days) as the response variable and germination timing (transplant date coded as continuous), species, their interaction, transplant height, bench, and population as fixed effects. Marginal means of the relationship between germination timing and time to first bud were estimated for each species and are reported along with standard errors (SE), and 95% confidence intervals (CI) used to evaluate significance. Confidence intervals in bold are significant.

| Species | Germination Timing | SE | CI |
|---|---|---|---|
| STDR | -0.69 | 0.05 | **-0.78 – -0.60** |
| STBR | -0.62 | 0.04 | **-0.71 – -0.54** |
| STTO | -0.94 | 0.11 | **-1.15 – -0.73** |
| STDI | -0.60 | 0.04 | **-0.69 – -0.52** |
| STPO | -0.50 | 0.05 | **-0.60 – -0.40** |
| STIN | -0.30 | 0.02 | **-0.34 – -0.26** |
| STGL | -0.56 | 0.06 | **-0.67 – -0.45** |
| CAAN | -0.49 | 0.03 | **-0.55 – -0.43** |
| CACO | -0.49 | 0.04 | **-0.57 – -0.42** |
| CAIN | -0.43 | 0.03 | **-0.49 – -0.36** |

**Table S8.** Results from the linear model with first bud date as the response variable and germination timing (transplant date coded as continuous), species, their interaction, transplant height, bench, and population as fixed effects. First bud date was calculated as days from September 1 to first bud for all individuals. Marginal means of the relationship between germination timing and first bud date were estimated for each species and are reported along with standard errors (SE), and 95% confidence intervals (CI) used to evaluate significance. Confidence intervals in bold are significant.

| Species | Germination Timing | SE | CI |
|---|---|---|---|
| STDR | 0.31 | 0.05 | **0.21 – 0.40** |
| STBR | 0.38 | 0.04 | **0.29 – 0.46** |
| STTO | 0.06 | 0.11 | -0.15 – 0.27 |
| STDI | 0.40 | 0.04 | **0.31 – 0.48** |
| STPO | 0.50 | 0.05 | **0.40 – 0.60** |
| STIN | 0.70 | 0.02 | **0.66 – 0.74** |
| STGL | 0.44 | 0.06 | **0.33 – 0.55** |
| CAAN | 0.51 | 0.03 | **0.45 – 0.57** |
| CACO | 0.51 | 0.04 | **0.43 – 0.58** |
| CAIN | 0.57 | 0.03 | **0.51 – 0.64** |

**Table S9.** Results from the linear model with size (height) at first bud as the response variable and germination timing (transplant date coded as continuous), species, their interaction, transplant height, bench, and population as fixed effects. Marginal means of the relationship between germination timing and height at first bud were estimated for each species and are reported along with 95% confidence intervals (CI), used to evaluate significance, and standard errors (SE). Confidence intervals in bold are significant.

| Species | Germination Timing | SE | CI |
|---|---|---|---|
| STDR | -0.01 | 0.01 | -0.02 – 0.01 |
| STBR | -0.02 | 0.01 | **-0.04 – -0.01** |
| STTO | 0.04 | 0.02 | -0.00005 - 0.08 |
| STDI | -0.10 | 0.01 | **-0.11 – -0.08** |
| STPO | -0.03 | 0.01 | **-0.05 – -0.01** |
| STIN | -0.02 | 0.01 | **-0.03 – -0.01** |
| STGL | 0.0006 | 0.01 | -0.02 – 0.02 |
| CAAN | 0.002 | 0.01 | -0.02 – 0.03 |
| CACO | 0.004 | 0.01 | -0.02 – 0.03 |
| CAIN | 0.001 | 0.01 | -0.02 – 0.02 |

**Table S10.** Results from the generalized linear model with probability of flowering as the response variable and germination timing (transplant date coded as continuous), species, their interaction, transplant height, bench, and population as fixed effects. Marginal means of the relationship between germination timing and probability of flowering were estimated for each species and are reported along with 95% confidence intervals (CI), used to evaluate significance, and standard errors (SE). Confidence intervals in bold are significant.

| Species | Germination Timing | SE | CI |
|---|---|---|---|
| STDR | 0.06 | 0.01 | **0.03 – 0.08** |
| STBR | -0.02 | 0.004 | **-0.02 – -0.01** |
| STTO | -0.03 | 0.01 | **-0.04 – -0.01** |
| STDI | -0.0002 | 0.01 | -0.1 – 0.01 |
| STPO | -0.02 | 0.004 | **-0.03 – -0.01** |
| STIN | -0.01 | 0.005 | -0.02 – 0.003 |
| STGL | 0.01 | 0.01 | -0.01 – 0.03 |
| CAAN | -0.02 | 0.01 | **-0.03 – -0.01** |
| CACO | -0.01 | 0.01 | -0.02 – 0.01 |
| CAIN | -0.01 | 0.003 | **-0.01 – -0.001** |

**Table S11.** Results from the generalized linear model with number of seeds as the response variable and germination timing (transplant date coded as continuous), species, their interaction, transplant height, bench, and population as fixed effects. Marginal means of the relationship between germination timing and number of seed were estimated for each species and are reported along with 95% confidence intervals (CI), used to evaluate significance, and standard errors (SE). Confidence intervals in bold are significant.

| Species | Germination Timing | SE | CI |
|---|---|---|---|
| STDR | -0.004 | 0.004 | -0.01 – 0.004 |
| STBR | -0.01 | 0.005 | **-0.02 – -0.002** |
| STTO | 0.01 | 0.01 | -0.01 – 0.03 |
| STDI | -0.03 | 0.005 | **-0.04 – -0.02** |
| STPO | 0.01 | 0.01 | -0.01 – 0.03 |
| STIN | -0.02 | 0.002 | **-0.03 – -0.02** |
| STGL | -0.006 | 0.006 | -0.02 – 0.004 |
| CAAN | -0.01 | 0.003 | **-0.01 – -0.003** |
| CACO | -0.01 | 0.005 | **-0.02 – -0.001** |
| CAIN | -0.02 | 0.01 | **-0.03 – -0.01** |

**Table S12.** Results from the linear model with seed mass as the response variable and germination timing (transplant date coded as continuous), species, their interaction, transplant height, bench, and population as fixed effects. Marginal means of the relationship between germination timing and seed mass were estimated for each species and are reported along with 95% confidence intervals (CI), used to evaluate significance, and standard errors (SE). Confidence intervals in bold are significant.

| Species | Germination Timing | SE | CI |
|---|---|---|---|
| STDR | -0.000008 | 0.000007 | -0.00002 – 0.000005 |
| STBR | -0.00001 | 0.000008 | -0.00003 – 0.000002 |
| STTO | 0.000005 | 0.00002 | -0.00003 – 0.00004 |
| STDI | -0.00002 | 0.000007 | **-0.00003 – -0.000009** |
| STPO | -0.0000002 | 0.000009 | -0.00002 – 0.00002 |
| STIN | -0.00004 | 0.000003 | **-0.00005 – -0.00003** |
| STGL | -0.00002 | 0.000009 | -0.00003 – 0.000003 |
| CAAN | -0.00001 | 0.000005 | **-0.00002 – -0.000002** |
| CACO | -0.00001 | 0.000007 | **-0.00003 – -0.000001** |
| CAIN | -0.000009 | 0.000006 | -0.00002 – 0.000003 |

**Table S13.** Results from the generalized linear model with first year fitness as the response variable and germination timing (transplant date coded as continuous), species, their interaction, transplant height, bench, and population as fixed effects. First year fitness was calculated as the probability of flowering multiplied by the number of seeds produced. Marginal means of the relationship between germination timing and first year fitness were estimated for each species and are reported along with 95% confidence intervals (CI), used to evaluate significance, and standard errors (SE). Confidence intervals in bold are significant.

| Species | Germination Timing | SE | CI |
|---|---|---|---|
| STDR | 0.001 | 0.005 | -0.01 – 0.01 |
| STBR | -0.02 | 0.004 | **-0.03 – -0.02** |
| STTO | -0.02 | 0.01 | **-0.03 – -0.01** |
| STDI | -0.03 | 0.005 | **-0.04 – -0.01** |
| STPO | -0.01 | 0.01 | -0.02 – 0.01 |
| STIN | -0.02 | 0.003 | **-0.03 – -0.02** |
| STGL | -0.004 | 0.007 | -0.02 – 0.01 |
| CAAN | -0.02 | 0.003 | **-0.02 – -0.01** |
| CACO | -0.01 | 0.005 | **-0.02 – -0.003** |
| CAIN | -0.02 | 0.004 | **-0.03 – -0.01** |

**Table S14.** Results of structural equation models testing for direct effects of germination timing (cohort) on fitness (seed count) and indirect effects through phenology, amount of time to first bud (days). Time to first bud was calculated as the number of days between individual transplant date and date of first bud. Numbers are standardized partial regression coefficients for direct effects along with calculations of indirect and total effects of each predictor variable (From) to each response variable (To). Direct effects in bold are significant with the number of * corresponding to significance level, * < 0.05, ** < 0.01, *** < 0.0001.

| **STDR** | | | | | |
|---|---|---|---|---|---|
| To | From | Direct | Indirect | Total | $R^2$ |
| Time to first bud | Cohort | **-0.8677*** ** | | -0.8677 | 0.83 |
| | Transplant height | -0.009 | | -0.009 | |
| Seed counts | Cohort | **-0.6310009*** ** | 0.6505763 | 0.0195754 | 0.62 |
| | Transplant height | **-0.1735114*** | 0.006747939 | -0.1667635 | |
| | Time to first bud | **-0.749771*** ** | | -0.749771 | |
| **STBR** | | | | | |
| To | From | Direct | Indirect | Total | $R^2$ |
| Time to first bud | Cohort | **-0.8926*** ** | | -0.8926 | 0.81 |
| | Transplant height | **-0.1034*** | | -0.1034 | |
| Seed counts | Cohort | **-0.1629701*** ** | 0.1330678 | -0.0299023 | 0.55 |
| | Transplant height | -0.01023675 | 0.01541476 | 0.00517801 | |
| | Time to first bud | **-0.1490789*** ** | | -0.1490789 | |
| **STTO** | | | | | |
| To | From | Direct | Indirect | Total | $R^2$ |
| Time to first bud | Cohort | **-0.8445*** ** | | -0.8445 | 0.72 |
| | Transplant height | 0.0758 | | 0.0758 | |

| | | | | | |
|---|---|---|---|---|---|
| Seed counts | Cohort | **-0.9947621**** | 1.183683 | 0.1889209 | 0.75 |
| | Transplant height | -0.02758675 | -0.1062442 | -0.133831 | |
| | Time to first bud | **-1.401638***** | | -1.401638 | |
| **STDI** | | | | | |
| To | From | Direct | Indirect | Total | $R^2$ |
| Time to first bud | Cohort | **-0.9615***** | | -0.9615 | 0.83 |
| | Transplant height | **-0.1558**** | | -0.1558 | |
| Seed counts | Cohort | **-0.8792721***** | 0.5237735 | -0.3554986 | 0.62 |
| | Transplant height | -0.02546689 | 0.08487146 | 0.05940457 | |
| | Time to first bud | **-0.5447462***** | | -0.5447462 | |
| **STPO** | | | | | |
| To | From | Direct | Indirect | Total | $R^2$ |
| Time to first bud | Cohort | **-0.689***** | | **-0.689***** | 0.55 |
| | Transplant height | **-0.2227*** | | **-0.2227*** | |
| Seed counts | Cohort | -0.008334085 | 0.006900514 | -0.001433571 | 0.15 |
| | Transplant height | 0.005477254 | 0.002230398 | 0.007707652 | |
| | Time to first bud | -0.01001526 | | -0.01001526 | |
| **STIN** | | | | | |
| To | From | Direct | Indirect | Total | $R^2$ |
| Time to first bud | Cohort | **-0.756***** | | **-0.756***** | 0.62 |
| | Transplant height | **-0.1882***** | | **-0.1882***** | |
| Seed counts | Cohort | **-0.4225315***** | 0.01878383 | -0.4037477 | 0.51 |
| | Transplant height | -0.02590242 | 0.004676081 | -0.02122634 | |
| | Time to first bud | -0.02484634 | | -0.02484634 | |
| **STGL** | | | | | |
| To | From | Direct | Indirect | Total | $R^2$ |

| | | | | | |
|---|---|---|---|---|---|
| Time to first bud | Cohort | **-0.7762*** ** | | -0.7762 | 0.59 |
| | Transplant height | 0.0127 | | 0.0127 | |
| Seed counts | Cohort | **-0.06218346** ** | 0.06165536 | -0.0005281 | 0.27 |
| | Transplant height | -0.02409811 | -0.00100879 | -0.0251069 | |
| | Time to first bud | **-0.07943231*** ** | | -0.07943231 | |
| **CAAN** | | | | | |
| To | From | Direct | Indirect | Total | $R^2$ |
| Time to first bud | Cohort | **-0.6148*** ** | | -0.6148 | 0.36 |
| | Transplant height | -0.082 | | -0.082 | |
| Seed counts | Cohort | **-0.6919082*** ** | 0.4223289 | -0.2695793 | 0.61 |
| | Transplant height | **0.1479089** ** | 0.05632883 | 0.2042377 | |
| | Time to first bud | **-0.686937*** ** | | -0.686937 | |
| **CACO** | | | | | |
| To | From | Direct | Indirect | Total | $R^2$ |
| Time to first bud | Cohort | **-0.8063*** ** | | -0.8063 | 0.64 |
| | Transplant height | 0.0264 | | 0.0264 | |
| Seed counts | Cohort | **-0.2633674*** ** | 0.1746934 | -0.088674 | 0.41 |
| | Transplant height | 0.003769832 | -0.005719837 | -0.001950005 | |
| | Time to first bud | **-0.2166605*** ** | | -0.2166605 | |
| **CAIN** | | | | | |
| To | From | Direct | Indirect | Total | $R^2$ |
| Time to first bud | Cohort | **-0.7461*** ** | | -0.7461 | 0.55 |
| | Transplant height | -0.0078 | | -0.0078 | |
| Seed counts | Cohort | **-0.4286871*** ** | 0.2778201 | -0.150867 | 0.46 |
| | Transplant height | 0.07935391 | 0.002904432 | 0.08225834 | |
| | Time to first bud | **-0.3723631*** ** | | -0.3723631 | |

**Table S15.** Results from phylogenetic generalized linear models used to account for phylogenetic relationships when evaluating relationships between time to first bud, flowering probability, number of seeds, and first year fitness slopes with 25-year average annual climate variables. Principal component analysis was used to reduce dimensionality of the climate variables (Table S6). Most climate variables were associated with PC1, so it was used as the predictor in these models. Coefficient estimates are reported for the Intercept and PC1 terms from the models along with standard error (SE) and the p-value associated with the PC1 term. $R^2$ values are also reported. The p-values in bold are significant.

| Response | Intercept | PC1 | SE | P-value | $R^2$ |
|---|---|---|---|---|---|
| Time to first bud | -0.57 | -0.074 | 0.018 | **0.003853** | 0.63 |
| Flowering Probability | -0.01 | -0.001 | 0.005 | 0.9221 | 0.001 |
| Number of Seeds | -0.01 | 0.005 | 0.002 | **0.01739** | 0.53 |
| Year 1 Fitness | -0.01 | 0.001 | 0.002 | 0.61093 | 0.03 |

**Table S16.** We used phylogenetic generalized linear models to test the hypothesis that species with narrower seasonal germination niches observed in our previous study (Worthy et al. 2023) might have greater fitness sensitivity when forced to experience seasonal conditions under which they normally would avoid germinating. Models evaluated the relationship between slopes of number of seeds produced and first year fitness against germination timing with slopes of relationships between germination proportion and rainfall onset date estimated in Worthy et al. (2023). Coefficient estimates are reported for the Intercept and slope terms from the models along with standard error (SE) and the p-value associated with the predictor term. $R^2$ values are also reported.

| Response | Intercept | Slope | SE | P-value | $R^2$ |
|---|---|---|---|---|---|
| Number of Seeds | -0.004 | 0.019 | 0.03 | 0.50 | 0.06 |
| Year 1 Fitness | -0.018 | -0.009 | 0.02 | 0.71 | 0.02 |

**Table S17.** We used phylogenetic generalized linear models to test the hypothesis that plasticity of flowering phenology to germination timing is related to stability of fitness to germination timing. Models evaluated the relationship between slopes of time to first bud and germination timing and three fitness components, slopes of number of seeds, flowering probability, and first year fitness with germination timing. Coefficient estimates are reported for the Intercept and slope terms from the models along with standard error (SE) and the p-value associated with the predictor term. $R^2$ values are also reported.

| Predictor | Intercept | Slope | SE | P-value | $R^2$ |
|---|---|---|---|---|---|
| Number of Seeds | -0.61 | -7.69 | 3.61 | 0.06581 | 0.36 |
| Flowering Probability | -0.54 | -0.59 | 2.08 | 0.78262 | 0.01 |
| Year 1 Fitness | -0.60 | -3.81 | 5.29 | 0.49116 | 0.06 |

**Table S18.** Principal component analysis of average climate for 1991-2015 for species' locations taken from herbarium records from the Consortium of California Herbaria. Climate variables include climate water deficit (CWD), precipitation (PPT), minimum and maximum temperature (Tmin, Tmax, respectively), as well as inter-annual variability in precipitation (coefficient of variation, PPT_CV) and temperature (standard deviation of Tmin and Tmax, Tmin_SD and Tmax_SD, respectively). The loadings of each climate variable on the first two principal component axes (PCs) along with the cumulative proportion of variation explained by the PCs are presented. The first two PCs are illustrated for each species in Figure S1. Data source was Flint and Flint, (2014).

| **Variable** | **PC1** | **PC2** |
|---|---|---|
| CWD | -0.509 | -0.001 |
| PPT | 0.489 | 0.089 |
| PPT_CV | -0.418 | -0.155 |
| Tmin | -0.371 | 0.067 |
| Tmin_SD | -0.067 | 0.691 |
| Tmax | -0.461 | -0.079 |
| Tmax_SD | -0.101 | 0.693 |
| Cumulative Proportion | 0.519 | 0.738 |

**Figure S1.** Location of populations used in this study within the species' range-wide climate space based on herbarium records from the Consortium of California Herbaria. A principal component analysis including all species was performed on climate variables from 1991-2015 (Table S18). Principal component scores were subset for each species and the first two principal components (PCs) are illustrated on the x and y axes, respectively. Climate variables included climate water deficit, precipitation, minimum and maximum temperature, as well as inter-annual variability in precipitation (coefficient of variation) and temperature (standard deviation). Climate data was sourced from the California Basic Characterization Model (Flint and Flint 2014). Population(s) included in this study are shown as a black triangle. For STTO, points are colored by elevation since this species spans a wide elevational range, but our study population is from low elevation (379 m). Panels are ordered by species' phylogenetic relationships.

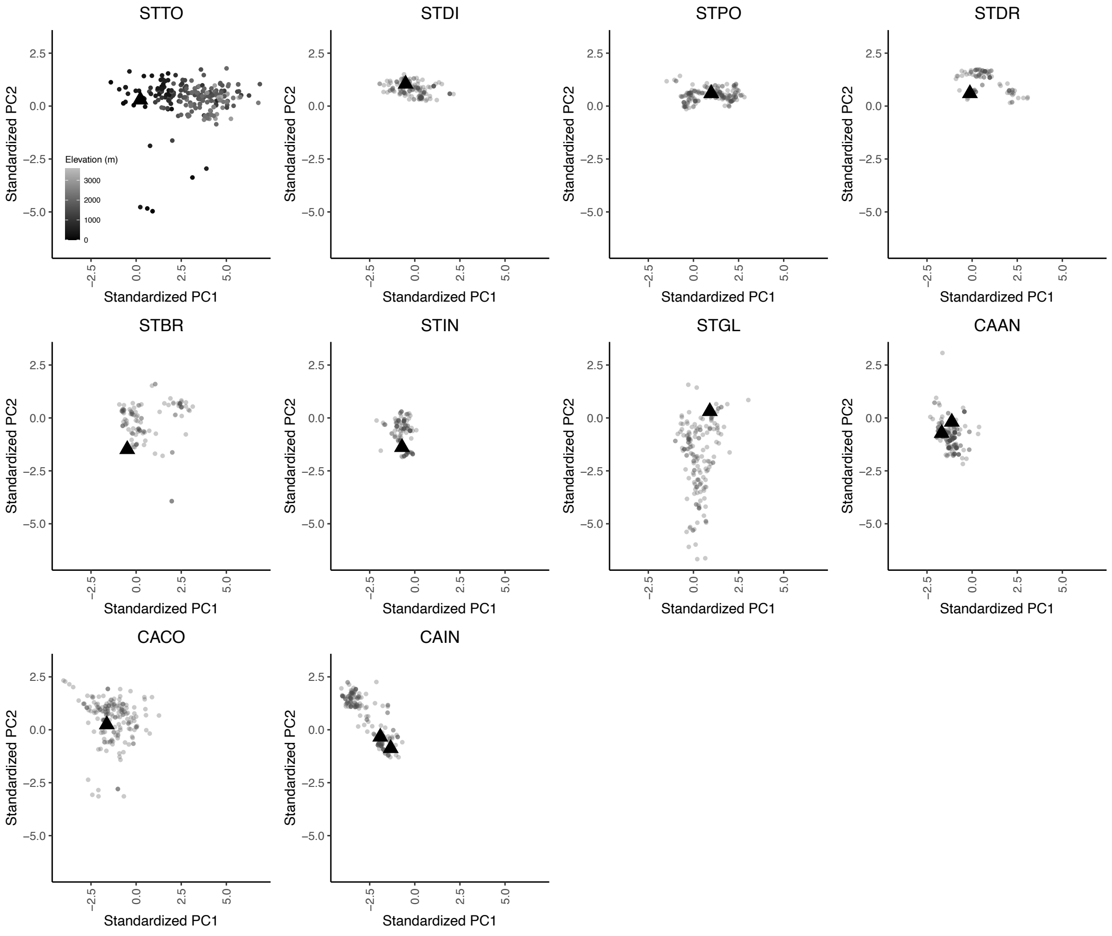

**Figure S2.** Seasonal conditions experienced differed for individuals in each of the germination timing cohorts. **(A)** The amount of chill portion and photothermal units accumulated by individuals decreased as the timing of rainfall and subsequent germination shifted later in the season. **(B)** Day length individuals in each of the germination timing cohorts experienced during the study. Chill portions and daylengths were calculated using the chillR package (Luedeling et al. 2023) for the location of the screenhouse. Photothermal units were calculated according to Burghardt et al. (2015) such that each germination timing cohort accumulated units on the basis of hourly temperatures and day lengths with a base temperature rate of 4°C. Dates on the x-axis correspond to transplant dates of the eight germination timing cohorts along with 13-Jun, the end date of the experiment.

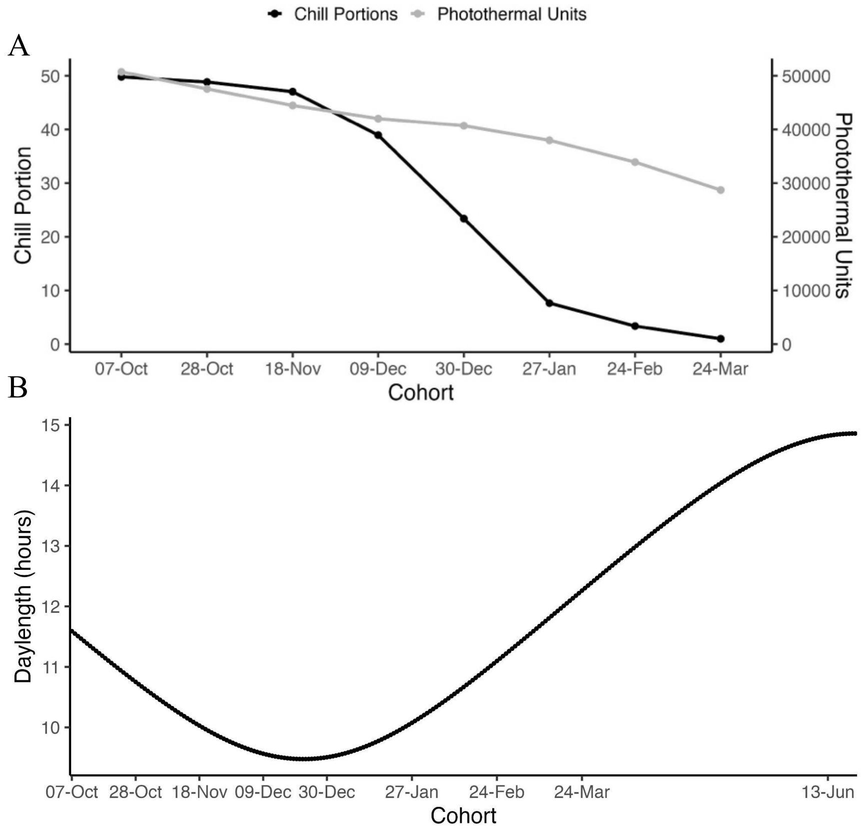

**Figure S3.** Relationships between first bud date and germination timing. First bud date was calculated as days from September 1 to first bud for all individuals. Regression lines represent significant, positive relationships where later germinating individuals produced their first bud later in the season, i.e., more days since September 1. Points represent observed days from September 1 for individuals of each species in each germination timing cohort. Panels are ordered by species' phylogenetic relationships.

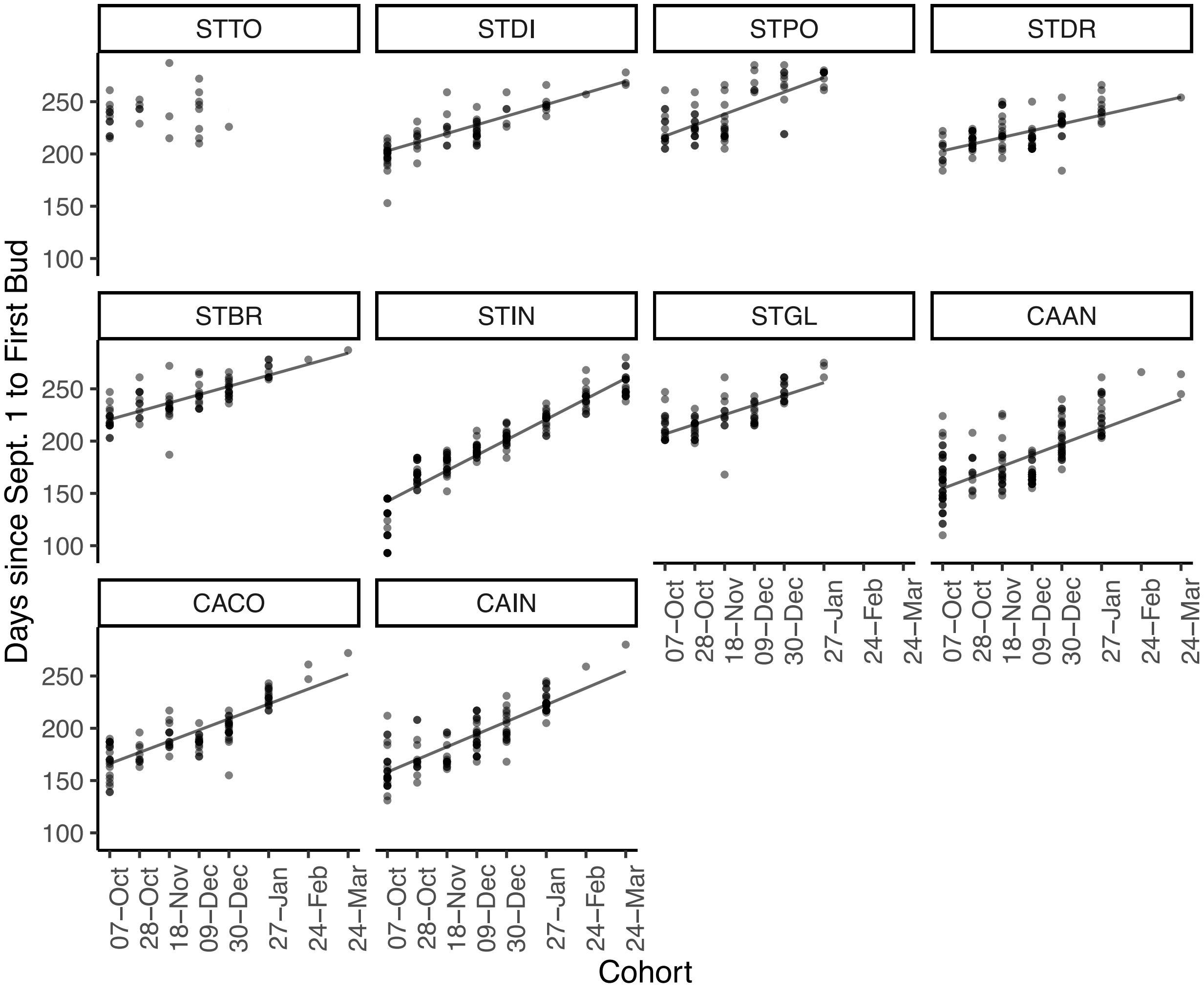

**Figure S4.** Relationships between size (height, cm) at first bud and germination timing. Regression lines represent significant, negative relationships where later germinating individuals were shorter when budding. Points represent the height at first bud for individuals of each species in each cohort. Panels are ordered by species' phylogenetic relationships.

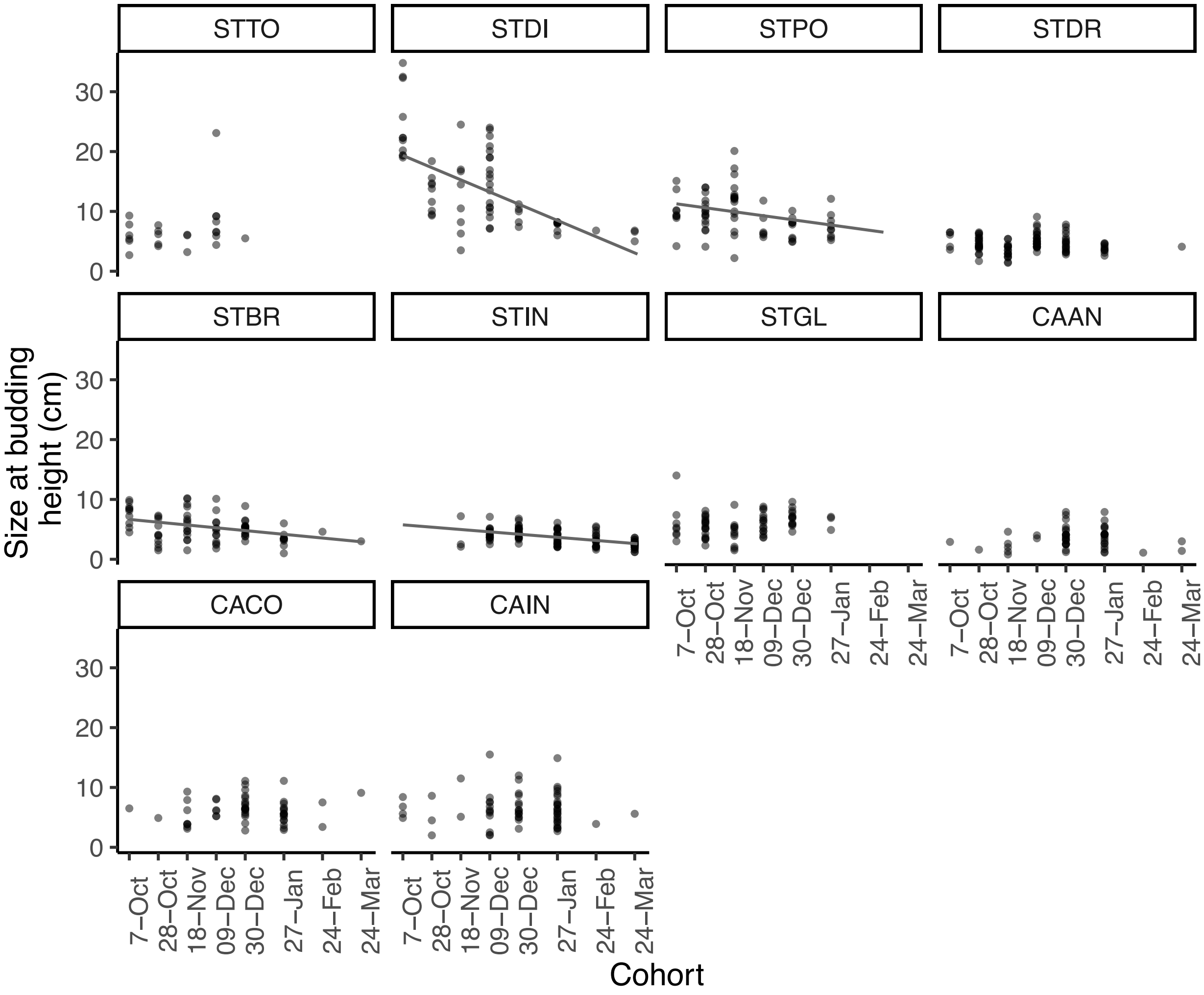

**Figure S5.** Proportions of individuals of each species in each cohort that fall into three categories: flowered during the experiment (black), lived until the end of the experiment but never flowered (dark gray), and died before reaching the first reproductive stage, budding (light gray). If bars are absent, no individuals germinated for these germination timing cohorts. Panels are ordered by species' phylogenetic relationships.

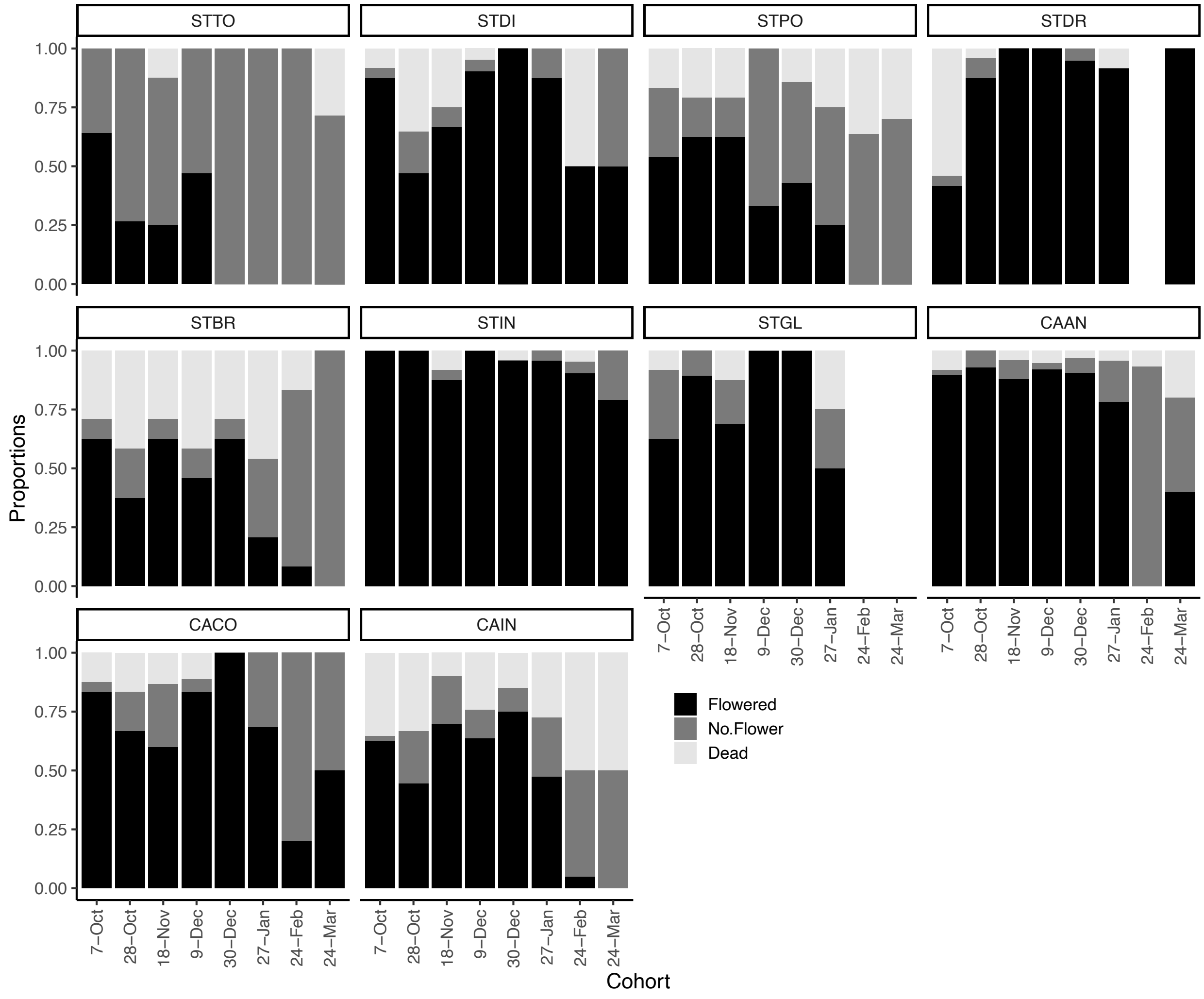

**Figure S6.** Relationships between seed mass (g) and germination timing. Regression lines represent significant, negative relationships where individuals that germinated later in the season had lower total seed mass. Points represent the seed mass for individuals of each species in each cohort. Panels are ordered by species' phylogenetic relationships.

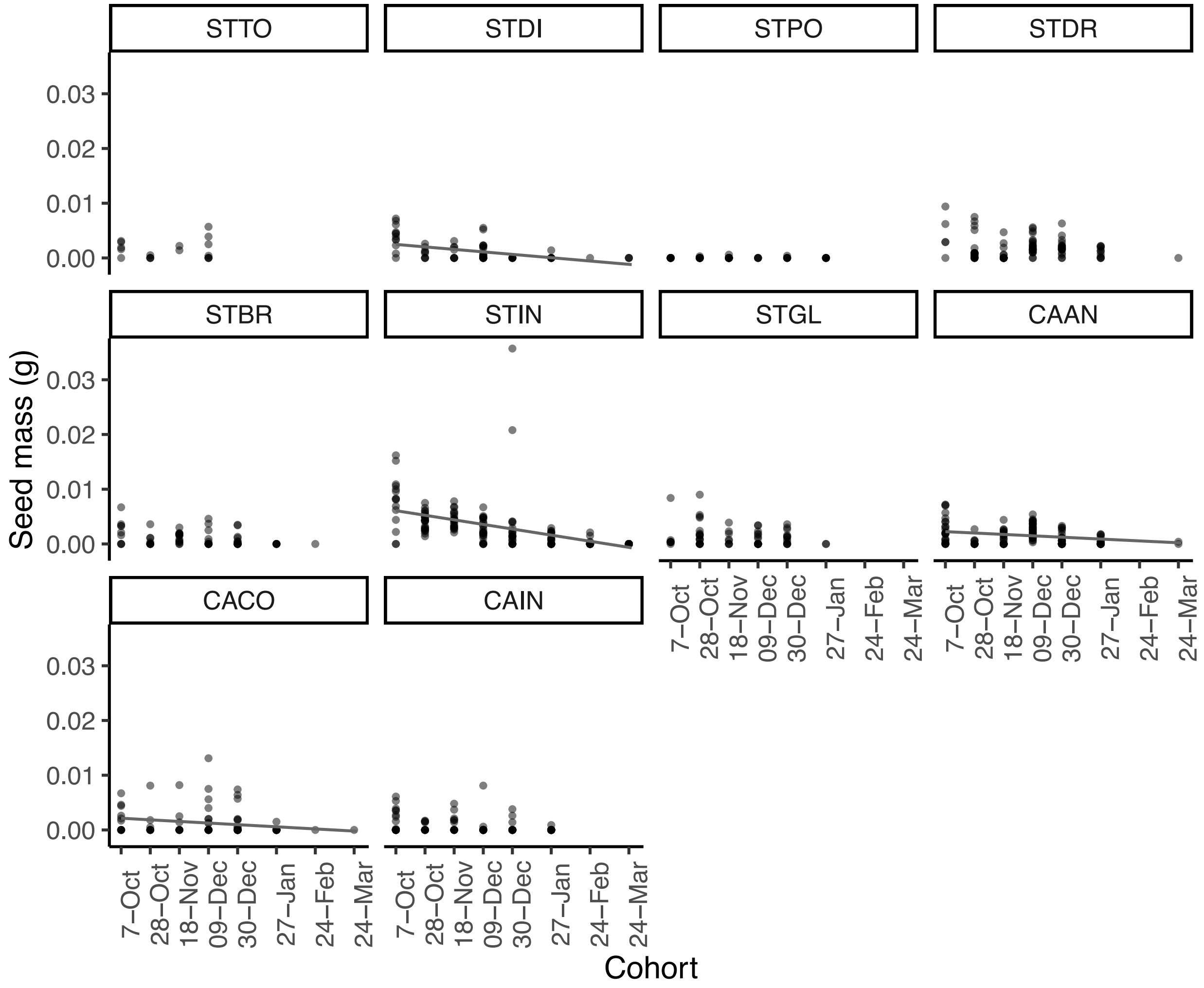

**Figure S7.** Relationships between first year fitness and germination timing. First year fitness was calculated as the probability of flowering multiplied by the number of seeds produced. Regression lines represent significant, negative relationships where later germinating individuals had lower first year fitness. Points represent first year fitness for individuals of each species in each germination timing cohort. Individuals with zero first year fitness flowered but produced zero fruits (n = 389) or did not flower (n = 503). Panels are ordered by species' phylogenetic relationships.

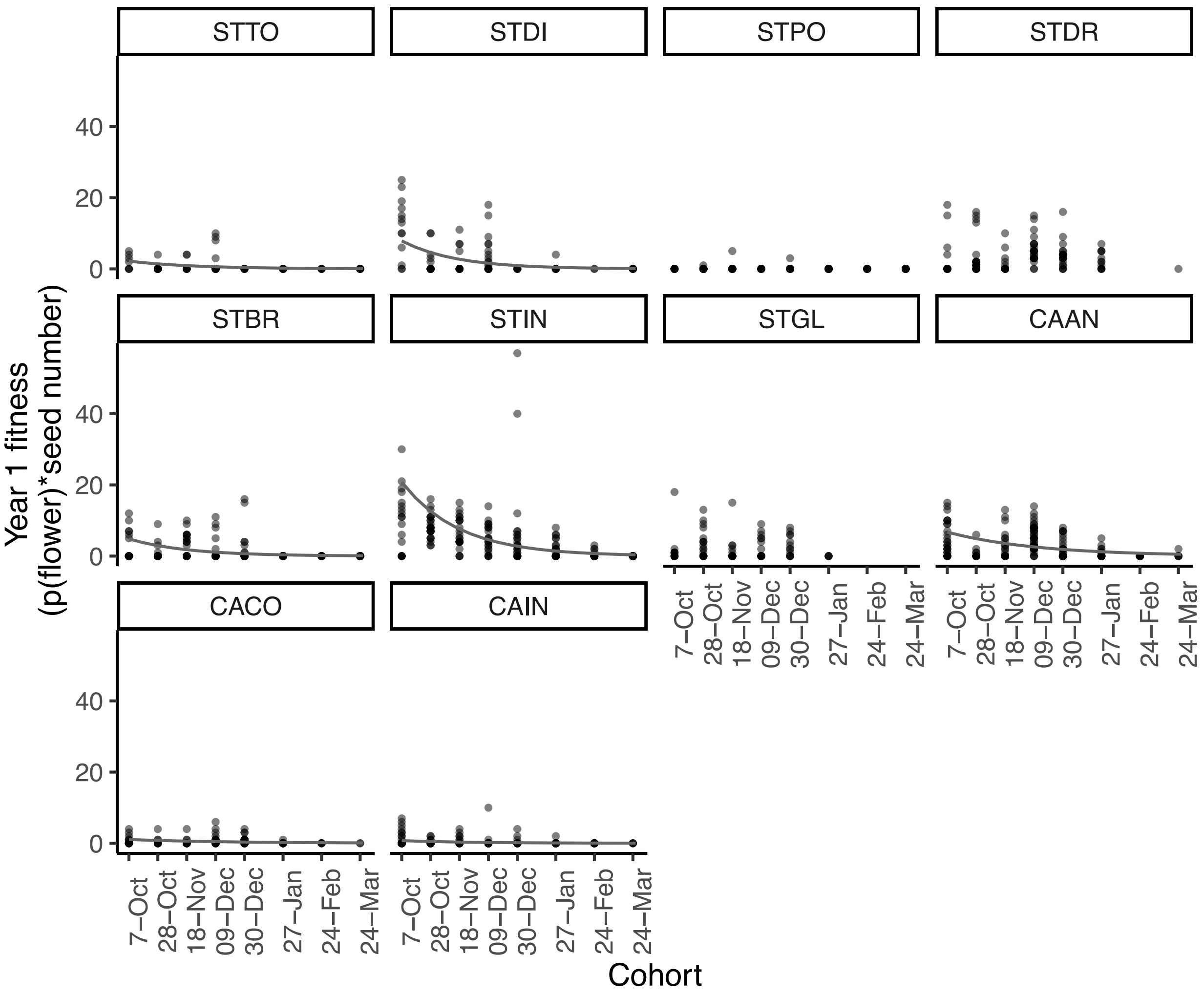

**Figure S8.** Slopes of relationships between first bud date, probability of flowering, total seed mass, first year fitness, and germination timing displayed across the phylogeny. First bud date was calculated as days from September 1 to first bud for all individuals. First year fitness was calculated as the probability of flowering multiplied by the number of seeds produced. Phylogenetic signal of these relationships was evaluated with Blomberg's K: first bud date (K = 0.18, p = 0.15); probability of flowering (K = 0.97, p = 0.86); total seed mass (K = 0.77, p = 0.39), and first year fitness (K = 0.68, p = 0.99).

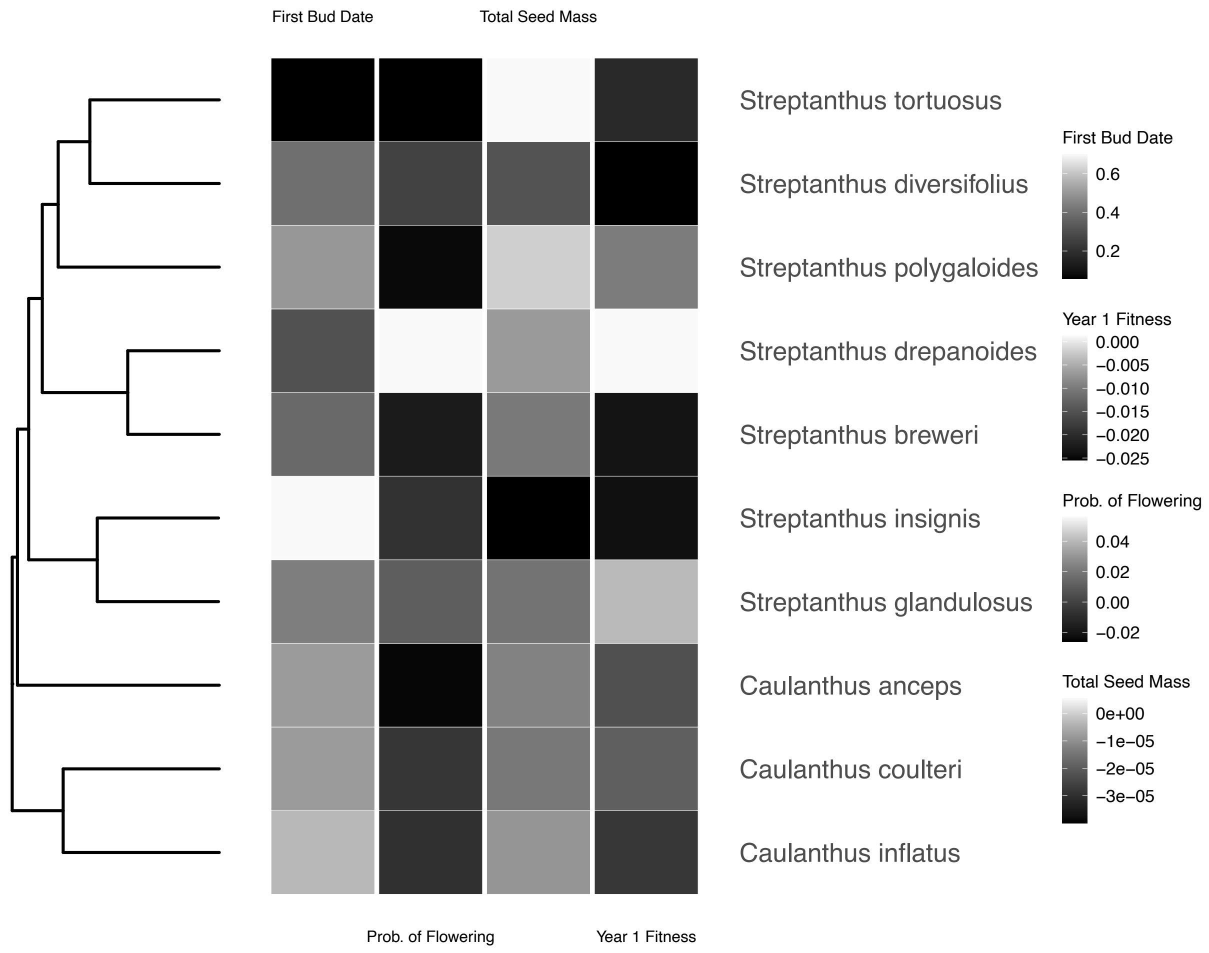

**Figure S9.** Relationships between **(A)** slopes of number of seeds produced against germination timing and **(B)** slopes of first year fitness against germination timing with slopes of relationships between germination proportion and germination timing from Worthy et al. (2023). Relationships were evaluated using phylogenetic generalized linear models (Table S16). First year fitness was calculated as the probability of flowering multiplied by the number of seeds produced. Points represent observed values for each species.

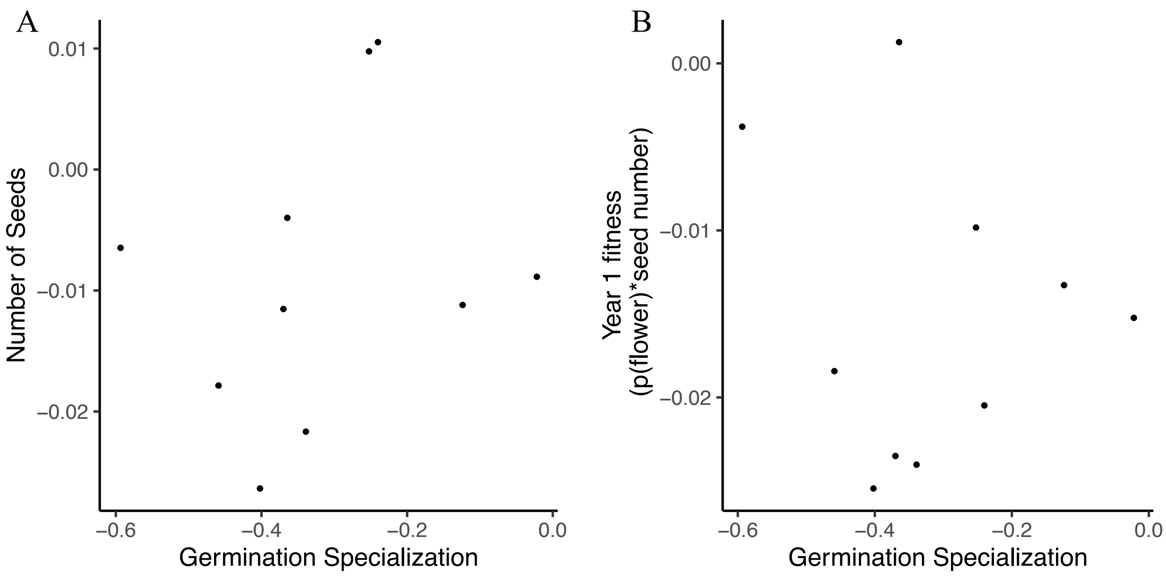

**Figure S10.** Relationships between slopes of time to first bud (days) against germination timing, with **(A)** slopes of number of seeds produced, **(B)** slopes of flowering probability, and **(C)** slopes of first year fitness against germination timing. Relationships were evaluated using phylogenetic generalized linear models (Table S17). First year fitness was calculated as the probability of flowering multiplied by the number of seeds produced. Points represent observed values for each species.

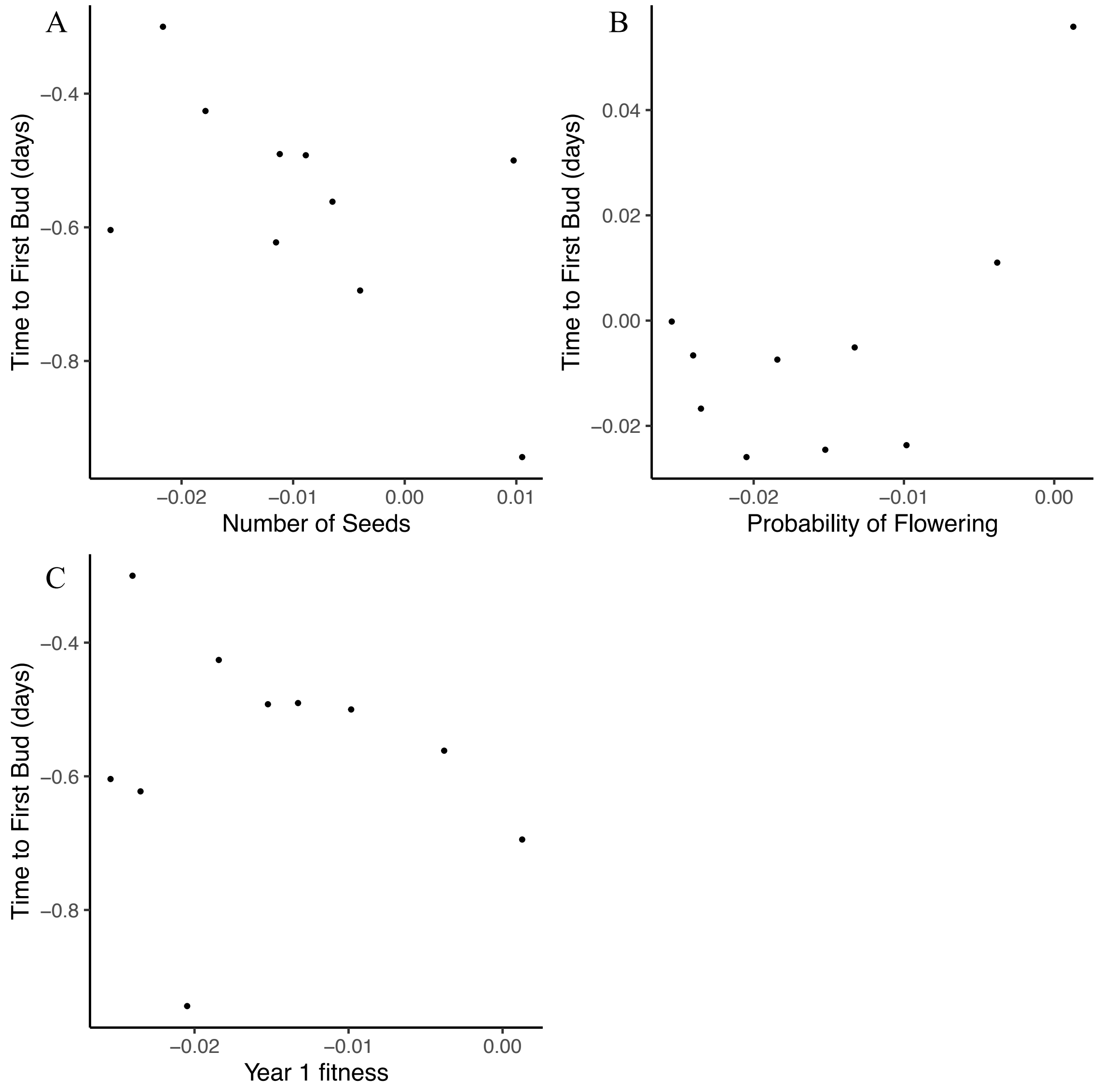
